# Detection of differentially abundant cell subpopulations discriminates biological states in scRNA-seq data

**DOI:** 10.1101/711929

**Authors:** Jun Zhao, Ariel Jaffe, Henry Li, Ofir Lindenbaum, Esen Sefik, Ruaidhrí Jackson, Xiuyuan Cheng, Richard Flavell, Yuval Kluger

## Abstract

Traditional cell clustering analysis used to compare the transcriptomic landscapes between two biological states in single cell RNA sequencing (scRNA-seq) is largely inadequate to functionally identify distinct and important differentially abundant (DA) subpopulations between groups. This problem is exacerbated further when using unsupervised clustering approaches where differences are not observed in clear cluster structure and therefore many important differences between two biological states go entirely unseen. Here, we develop DA-seq, a powerful unbiased, multi-scale algorithm that uniquely detects and decodes novel DA subpopulations not restricted to well separated clusters or known cell types. We apply DA-seq to several publicly available scRNA-seq datasets on various biological systems to detect differences between distinct phenotype in COVID-19 cases, melanomas subjected to immune checkpoint therapy, embryonic development and aging brain, as well as simulated data. Importantly, we find that DA-seq not only recovers the DA cell types as discovered in the original studies, but also reveals new DA subpopulations that were not described before. Analysis of these novel subpopulations yields new biological insights that would otherwise be neglected.

## Introduction

Profiling biological systems with single cell RNA sequencing (scRNA-seq) is an invaluable tool, as it enables experimentalists to measure the expression levels of all genes over thousands to millions of individual cells [1, 2]. A prevalent challenge in scRNA-seq analysis is comparing the transcriptomic profiles of cells from two biological states [3, 4]. Often, such comparison reveals subpopulations that are *differentially abundant* (DA). In DA subpopulations, the ratio between the number of cells from the two biological states differs significantly from the respective ratio in the overall data. Creation of a methodology to accurately resolve these differences have the potential to reveal significant novel observations currently hidden within datasets of major importance to human health such as COVID-19 and cancer immunotherapy. Generating new analytical pipelines with the power to decode all important information in scRNA-seq is a major challenge and impediment to fully unlocking these datasets for betterment of human disease and more generally biological systems.

A standard approach to detect DA subpopulations is by clustering the union of cells from both states. This step can be done in a completely unsupervised manner or by using known biomarkers. For each cluster, the proportion of cells from the two biological states is measured. A cluster in which these proportions significantly differ from the overall proportion in the data is considered differentially abundant. Often, a subset of differentially expressed genes that characterize each DA cluster is then identified. This approach was applied in the analysis of various biological systems, for example, to investigate immune response and mechanisms on patients with various diseases severity after viral infection [5, 6], to compare responders and non-responders to cancer treatment [7], and to study cell remodeling in inflammatory bowel disease [8]. A similar cluster-based method is ClusterMap [9], where the clustering step is applied separately to cells from the two states. Subsequently, the datasets are merged by matching similar clusters. Skinnider et al. [10] developed Augur, which employs machine learning to quantify *separability* of cells from two states within clusters. Comparing biological states through clustering is also related to differential compositional analysis, where biological states are compared via the proportion of predetermined cell types [11].

Clustering-based methods might be suboptimal, however, in cases where the subpopulations most responsive to the biological state do not fall into well-defined separate clusters. For example, DA subpopulations may be distributed among several adjacent clusters or, alternatively, encompass only a part of a cluster. Additionally, the clustering approach may fail for continuous processes where no clear cluster structure exists, such as cell cycles or certain developmental programs.

A different method for identifying DA subpopulations that does not rely on initial clustering was derived by Lun et. al. [12] for mass cytometry data. Their algorithm performs multiple local two sample tests for hyperspheres centered at randomly selected cells. The caveat of this approach is that the selected hyperspheres may only partially overlap with the DA subpopulations or fail to form localized regions. Accurate delineation of a DA subpopulation is essential for identifying the markers that differentiate it from its immediate neighboring cells as well as markers that separate it from the rest of the cells in the dataset.

Here, we develop DA-seq, a multiscale approach for detecting DA subpopulations. In contrast to clustering-based methods, DA-seq detects DA subpopulations that do not necessarily match predetermined clusters. For each cell, we compute a multiscale differential abundance score measure. These scores are based on the *k* nearest neighbors in the transcriptome space for various values of *k*. The motivation of multiscale analysis is that by employing a single scale, one may miss some of the DA subpopulations if the scale is too large, or detect spurious DA subpopulations if the scale is too small. We applied DA-seq to various scRNA-seq datasets from published works as well as simulated datasets. We show that DA-seq successfully recovers findings presented in the original works. More importantly, DA-seq reveals new DA cell subpopulations that were not reported before. Characterization of these novel subpopulations provided insights crucial to understanding the biological processes and mechanisms.

## Results

### The DA-seq algorithm

Here, we briefly outline the main four steps of the DA-seq algorithm (Fig. 1). As a first step, DA-seq computes for each cell a *score* based on the relative prevalence of cells from both biological states in the cell’s neighborhood. Importantly, this measure is computed for neighborhoods of different size, thus providing a *multiscale* measure of differential abundance for each cell. In the second step, the multiscale measure is merged into a single *DA measure* as quantity of differential abundance. This step is done by training a logistic regression classifier to predict the biological state of each cell. The associated prediction probability is then used as a DA measure of how much a cell’s neighborhood is dominated by cells from one of the biological states. In the third step, DA-seq clusters the cells whose DA measure is above or below a certain threshold into localized regions based on gene expression profiles. The cells in each region represent cell subpopulations with a significant change in abundance between biological states. In the final step, DA-seq selects genes that distinguish a DA subpopulation from the rest of the cells in the data or cells from its immediate neighborhood. As detailed in the Methods section, for this task we employ a novel *l*_0_ based feature selection method via stochastic gates (STG) [13], thereby identifying the minimum number of genes that distinguish a DA subpopulation. However, standard differential expression methods may be applied here as well. The four steps of DA-seq are outlined in Algorithm 1 and illustrated in Fig. 1. All steps are described in detail in the Methods section.

**Figure 1:**
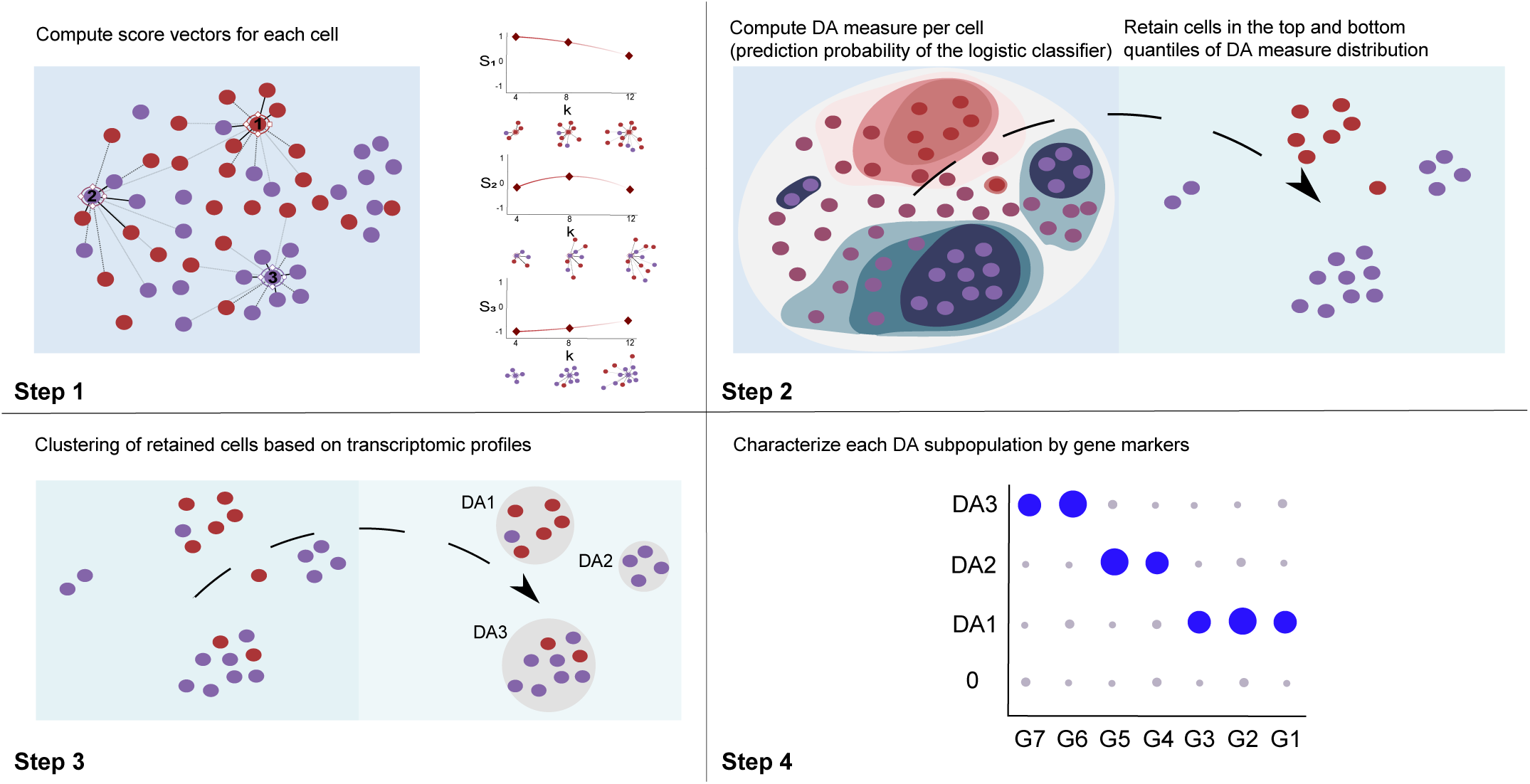
Schematic demonstration of DA-seq. Our method detects DA subpopulations by analyzing cells from two biological states, colored by red and purple, respectively. **Step 1:** Computing a *multiscale score* measure for each cell according to Eq. (6) for several values of *k* (e.g. *k* = 4, 8, 12). **Step 2:** Training a logistic classifier to predict the biological state of each cell based on the multiscale score (see Eq. (7)) to obtain a single *DA measure*. The algorithm retains only cells for which the output is above a threshold *τ*_*h*_ or below *τ*_*l*_ and hence may reside in DA subpopulations. **Step 3:** Clustering the cells retained in Step 2 to obtain contiguous and non-negligible DA subpopulations. These subpopulations are denoted *DA*1, *DA*2 and *DA*3. **Step 4:** Detect subsets of genes that characterize each of the DA subpopulations by applying standard differential expression analysis and/or feature selection via stochastic gates (STG). For example, the genes *G6* and *G7* characterize *DA*3.

To quantify the differential abundance for every detected DA subpopulation in Step 3, we define a *DA-score*. The form of this score is similar to the element-wise score of each individual cell as defined in Eq. (6). In addition, when adequate biological replicates are available, we provide *p*-values for DA subpopulations to assess reproducibility. More details regarding the calculation of the *DA-score* and *p*-value are given in Supplementary Note 1.

We applied DA-seq to publicly available scRNA-seq datasets from diverse biological systems [5, 7, 14, 15]. In the following sections, we present the output of steps 2, 3 and 4 of the DA-seq algorithm for datasets from [5, 7, 14]. We then compare the results to the findings in the original works, and validate our findings. Importantly, we show that DA-seq provides invaluable biological insights through the characterization of DA subpopulations that are not revealed by standard clustering-based approaches. Additional results on dataset from Ximerakis et al. [15] and simulated datasets can be found in Supplementary Information.

### Abundance of immune cell subsets in responsive vs. non-responsive melanoma patients

One of the goals of the Sade-Feldman et al. study [7] was to identify factors related to the success or failure of immune checkpoint therapy. To that end, 16,291 immune cells from 48 samples of melanoma patients treated with checkpoint inhibitors were profiled and analyzed. The tumor samples were classified as responders or non-responders based on radiologic assessments. The cells originating from responding tumors and non-responding tumors are labeled in the t-SNE plot of Fig. 2a. Comparisons between responders and non-responders yielded important biological insights.

**Figure 2:**
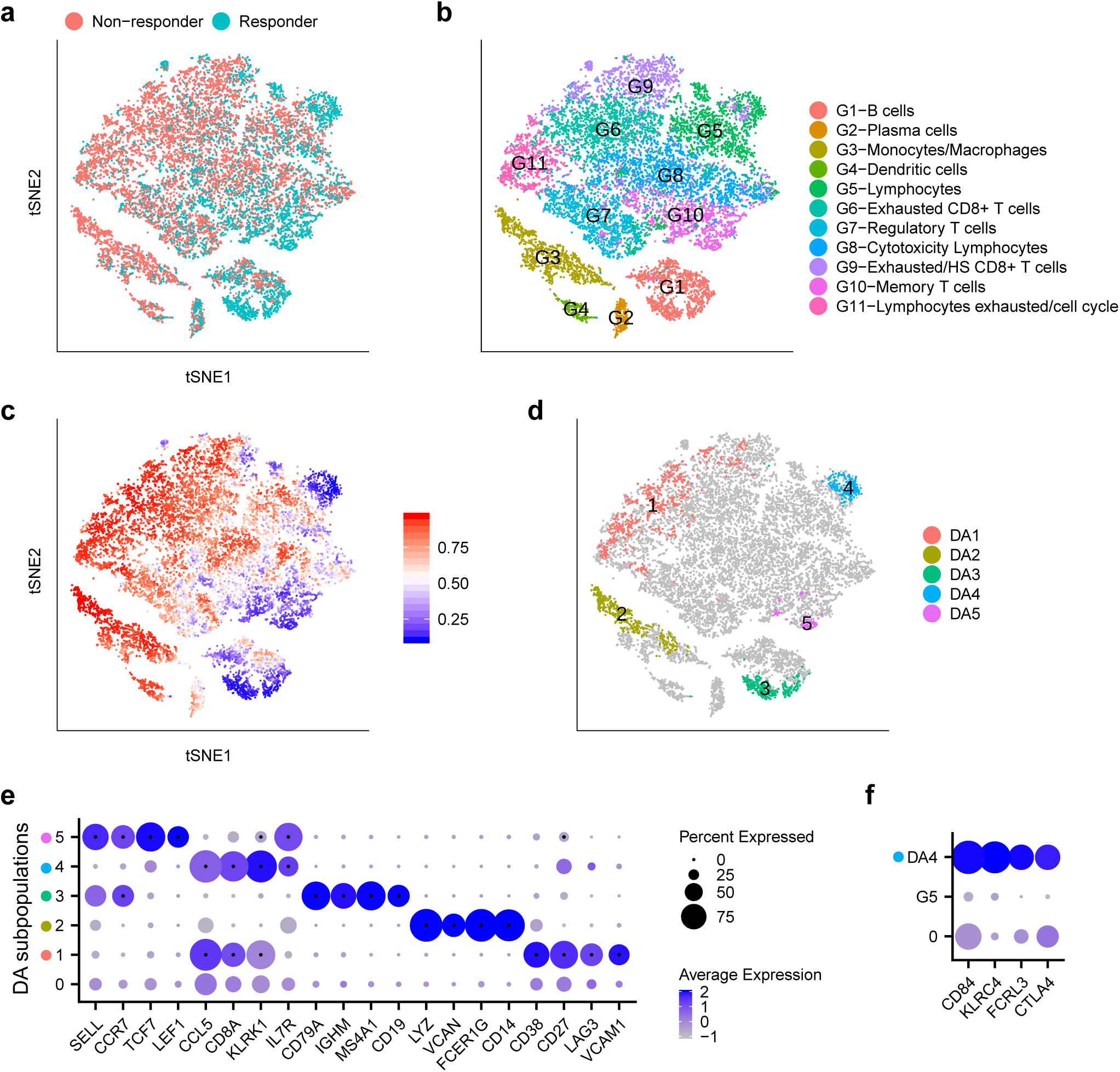
Immune cells from responding and non-responding melanoma patients treated with checkpoint therapy. **a-d** t-SNE embedding of 16,291 cells from [7]. **a** Cells colored by status of response to immune therapy. **b** Cells colored by cluster labels from [7]. **c** Cells colored by DA measure. Large (small) values indicate a high abundance of cells from the pool of non-responder (responder) samples. **d** Five distinct DA subpopulations obtained by clustering cells from the top 10% and bottom 5% quantiles (see Choice of Parameters) of the DA measure in **c. e** Dot plot for markers characterizing the five selected DA subpopulations. The color intensity of each dot corresponds to the average gene expression across all cells in the DA subpopulation excluding the cells with zero expression values. The lowest row in the plot corresponds to the non-DA cells (cells not included in any DA subpopulations). For each DA subpopulation (row), we mark the genes selected by STG with a black point in the center of the respective dots. **f** Dot plot for markers that distinguish *DA*4 and the complementary cells within *G*5.

Sade-Feldman et al. clustered the 16,291 immune cells into 11 distinct clusters (Fig. 2b). Subsequently, they computed the percentage of cells in each of the predefined clusters from responder and non-responder samples and compared the relative abundance between these two groups. Two clusters (*G*1, *G*10) were enriched in cells from the responder samples, and four clusters (*G*3, *G*4, *G*6, *G*11) were enriched in cells from non-responder samples. Finally, the authors composed a list of genes that were highly expressed in these six differentially abundant clusters.

Fig. 2c shows the intensity of the DA measure of each cell as computed in Step 2 of the algorithm, where higher values indicate an abundance of cells from non-responder samples relative to responder samples. Five DA cell subpopulations denoted *DA*1-*DA*5 (Fig. 2d, Supplementary Fig. S1a) were identified. In contrast to the method applied in [7], the DA subpopulations obtained by our approach are not constrained to any predefined clusters. Thus, there are some important differences between our findings and those of [7] despite some similarities. Five out of the six DA clusters described in [7] have partial overlaps with our DA subpopulations:

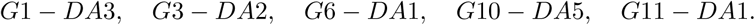

In [7], the clusters *G*11 and *G*6 are reported as two distinct DA clusters. In contrast, our DA cell subpopulation *DA*1 overlaps with both *G*6 and *G*11, as well as another cluster *G*9. We argue that identification of *G*6 and *G*11 as two separate DA clusters and the exclusion of *G*9 as potentially relevant for DA are artificial. Unifying the clusters of exhausted lymphocytes allows us to detect and transcriptionally characterize cell subpopulations within this union that are more specific to differences between responders and non-responders. We observe that DA subpopulations *DA*3, *DA*2 and *DA*5 partially overlap with *G*1, *G*3 and *G*10, respectively, but they are not identical; furthermore, subpopulation *DA*4 partially overlaps with cluster *G*5 which was not identified as a DA cluster. Unlike the predefined clusters that could bias the differential abundance analysis and result in diffused DA clusters, our method simply finds the most discriminative cells regardless of whether the underlying data has either a cluster structure or is a contiguous data cloud.

The cluster *G*4 (dendritic cells) which was reported in [7] as a DA cluster, was not detected by DA-seq as a DA subpopulation. We note, however, that this subpopulation is detected with a slight relaxation of the upper threshold *τ*_*h*_ in Step 2 (Supplementary Fig. S2a).

Finally, we identified markers that characterize the DA subpopulations by both standard differential expression approach implemented in *Seurat* [16, 17] *and our novel feature selection approach via STG (see Methods). A subset of the identified markers are shown by a dot plot in Fig. 2e. For the DA subpopulations DA*2-*DA*5, DA-seq detected similar lists of characteristic markers to their corresponding clusters in Fig. 2b.

Interestingly, the characteristic markers *LAG3* and *CD27* for subpopulation *DA*1 define an exhausted lymphocyte population [18, 19] covering three clusters associated with lymphocyte exhaustion. Notably, *VCAM1* was the most significant gene in *DA*1 (Supplementary Fig. S2b), which covers parts of clusters *G*6, *G*9 and *G*11. Although *VCAM1* was reported in [7], it was not among the salient markers of their analysis. Analyzing these clusters separately diminished the significance of *VCAM1* relative to other genes. *VCAM1* expression on a class of cells discovered by the DA approach is intriguing, as it is a critically important cell adhesion and costimulatory ligand in the immune system [20]. In addition, *VCAM1* has been implicated as having an important role in immune escape as has been studied in [21, 22, 23, 24, 25].

To distinguish subpopulation *DA*4 and its immediate neighborhood, we performed differential expression analysis comparing *DA*4 and cells in cluster *G*5 that are not within *DA*4. This uncovered the distinct transcriptional profile of *DA*4 (Fig. 2f). Intriguingly, the *CTLA*4 gene is highly expressed in *DA*4 (enriched in responders). Incidentally, this gene was reported as a marker for non-responders in other cell types from [7]. Clustering-based DA analysis failed to detect this DA subpopulation and thus, missed this important insight.

Compared with standard differential expression approaches that simply output individual genes in a univariate manner, STG provides a prediction score (see Methods) as a linear combination of its selected genes that best separate each DA subpopulation from the rest of the cells. The improved discrimination by STG compared to a univariate approach is demonstrated in Supplementary Fig. S2c,d for DA subpopulations *DA*4 and *DA*5.

To asses the stability of DA-seq results, the following cross-validation procedure was performed. We split the data randomly into two sets *s*1 and *s*2, such that each set contains half non-responder samples and half responder samples. To compare the two sets, the same t-SNE embedding as in Fig. 2 is used to show the response status (Supplementary Fig. S3a,e) and cluster label (Supplementary Fig. S3b,f) for each cell. Next, we applied DA-seq separately to each set. The DA measure for both sets is shown in Supplementary Fig. S3c,g. Seven DA subpopulations denoted as *s*1*DA*1-*s*1*DA*7, and six DA subpopulations denoted as *s*2*DA*1-*s*2*DA*6, were detected from *s*1 and *s*2, respectively (Supplementary Fig. S3d,h). The characteristic genes of DA subpopulations in *s*1 and *s*2 are shown in Supplementary Fig. S3i,j. We observe that most of the DA subpopulations detected in *s*1 share common characteristic genes with their counterparts in *s*2, as well as in the full dataset. The exact match between DA subpopulations in *s*1, *s*2 and the full dataset is shown in Supplementary Fig. S3k. We notice that, subpopulation *s*1*DA*4 does not overlap with subpopulations from *s*2 or the full dataset when we apply the same threshold parameters. But with relaxed *τ*_*h*_ on the full dataset, *s*1*DA*4 overlaps with *DA*3 in Supplementary Fig. S2a. Further, subpopulation *DA*5 in the full dataset overlaps with *s*1*DA*7 in *s*1, but does not overlap with any subpopulations in *s*2. This may indicate that this DA subpopulation only exists in a subset of patients, as reflected by the *p*-values computed for each subpopulation (Supplementary Fig. S1a).

### Differentiation patterns of early mouse dermal cells

We applied DA-seq to scRNA-seq data from a study on developing embryonic mouse skin [14]. Cells from dorsolateral skin were sequenced for two time points of embryonic development (days E13.5 and E14.5), each with two biological replicates (Fig. 3a). Dermal cells were selected for analysis by using the marker *Col1a1* to study hair follicle dermal condensate (DC) differentiation.

**Figure 3:**
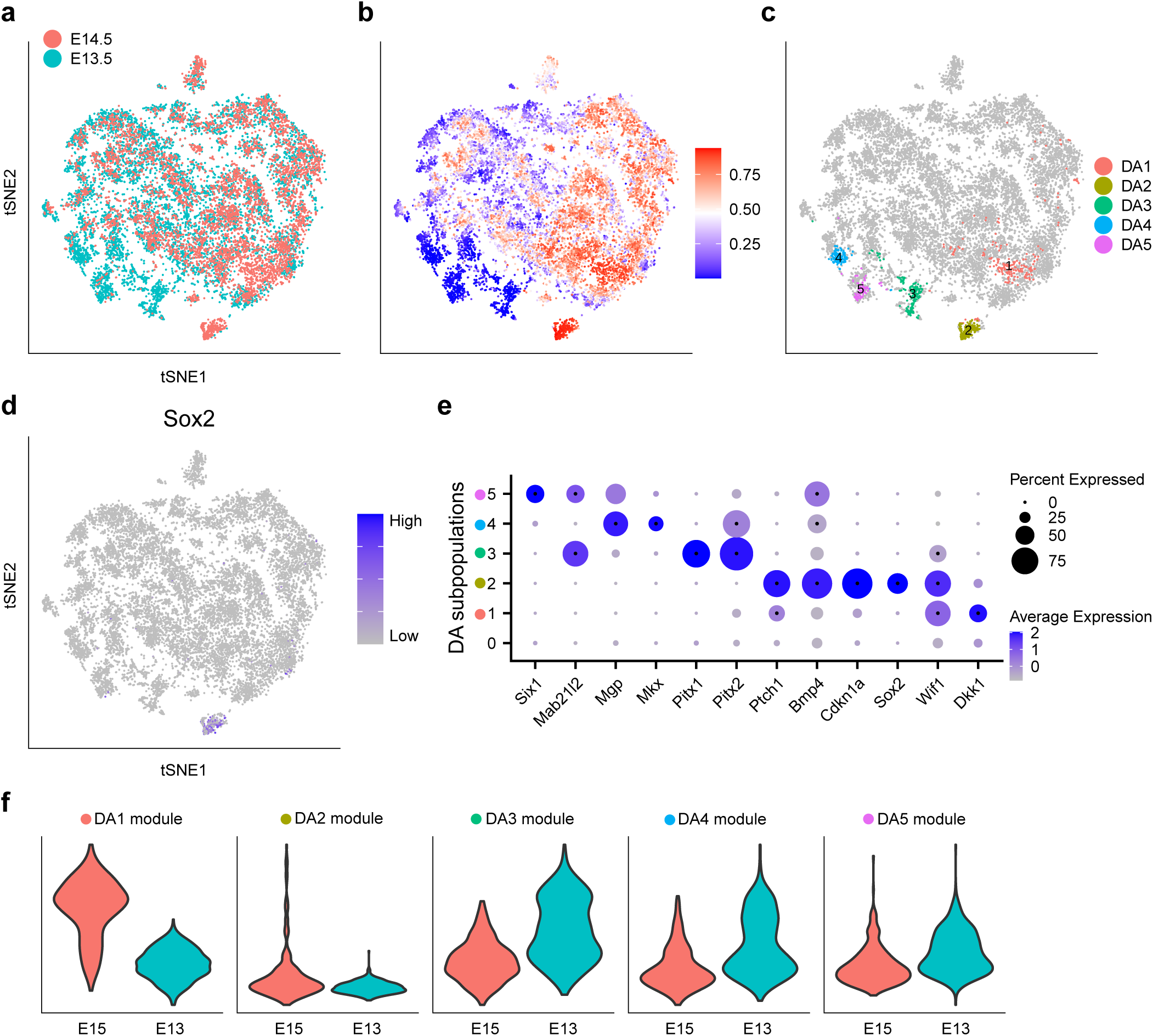
Comparing embryonic mouse dermal cells in embryonic day E13.5 and E14.5. **a-e** Show data from Gupta et al. [14]. **a-d** t-SNE embedding of 15,325 cells. **a** Embryonic day of each cell. **b** Cells colored by DA measure. Large (small) values indicate a high abundance of cells from E14.5 (E13.5). **c** Distinct DA subpopulations obtained by clustering cells from the top and bottom 3% quantiles of the DA measure in **b. d** Normalized *Sox2* gene expression. **e** Dot plot of several markers that characterize DA subpopulations. Details are as in Fig. 2**e. f** Validation on data from Fan et al. [26]. Violin plots comparing gene module scores between E15 and E13 samples in dermal cells of data from [26]. Gene modules are defined from DA subpopulations in **c**.

Gupta et al. [14] studied the transcriptional states of the cells by embedding them via diffusion map coordinates to capture the manifold structure of the scRNA-seq data. They then used the early DC marker *Sox2* to identify differentiated DC cells as well as the diffusion map dimension that corresponds to DC cell differentiation, which they called the DC-specific trajectory. By observing this trajectory, they found that although it contained cells from both E13.5 and E14.5, there were notably more E14.5 cells at the terminus representing differentiated DC cells.

In [14], the authors had prior knowledge that differentiated DC cells express *Sox2*. In contrast, DA-seq does not require prior knowledge. We obtained an unbiased comparison of dermal cells (Fig. 3a) between E13.5 and E14.5, which revealed the differentiated DC cell population discussed in [14]. Fig. 3b colors each cell according to its DA measure from Step 2 of DA-seq. Fig. 3c shows five DA subpopulations identified by DA-seq, denoted as *DA*1 − *DA*5. Fig. 3e shows the characteristic marker genes associated with these DA subpopulations. Due to lack of replicated samples in this dataset (two replicates for both E13.5 and E14.5), we did not compute a *p*-value. Instead, we computed the *DA-score* for these DA subpopulations for every possible pairwise comparison of these samples (Supplementary Fig. S1b). Specifically, E13.5 replicate 1 vs. E14.5 replicate 1, E13.5 replicate 1 vs. E14.5 replicate 2, E13.5 replicate 2 vs. E14.5 replicate 1, and E13.5 replicate 2 vs. E14.5 replicate 2.

Among the identified DA subpopulations, *DA*1 and *DA*2 are more abundant in E14.5. Subpopulation *DA*2 corresponds to the *Sox2*+ differentiated DC cells (Fig. 3d). Markers of *DA*2 (Fig. 3e) include other genes (*Cdkn1a,Bmp4, Ptch1*) known to be expressed in differentiated DC cells. Subpopulation *DA*1, characterized by the gene *Dkk1*, corresponds to a subpopulation that spatially surrounds the DC population. Although this subpopulation was acknowledged briefly in [14], the localization of *DA*1 in our analysis provides a method to interrogate the molecular mechanisms that regulate DC maturation and hair follicle development. Other characteristic markers of *DA*1 provide insights on more detailed biological functions of this peri-DC subpopulation. DA subpopulations *DA*3, *DA*4 and *DA*5 are more abundant in E13.5. Marker genes of these subpopulations (Fig. 3e) are associated with various developmental processes, potentially representing cell development or relocalization during early embryonic days.

To validate findings obtained by analyzing the data with DA-seq, we examined scRNA-seq data from another closely related study [26]. In [26], single cells isolated from the dorsal skin at embryonic days E13 and E15 were profiled. We defined gene signatures (see Methods) that are enriched in each of the five DA subpopulations detected in the data from Gupta et al. [14] shown in Fig. 3c. Gene modules scores (see Methods) for these signatures are computed and compared between E13 and E15 in dermal cells from Fan et al. [26]. The differences between the module score distributions of E15 versus E13 (Fig. 3f) are consistent with the enrichment of these signatures within the DA subpopulations in Fig. 3c.

### Severe and moderate COVID-19 patients have distinct immunological profiles

Coronavirus disease 2019 (COVID-19) is a current global pandemic of a novel virus. It is crucial to understand the immunological mechanisms related to disease severity. In [5], Chua et al. applied scRNA-seq on nasopharyngeal (nasopharyngeal or pooled nasopharyngeal/pharyngeal swabs (NSs)) samples from 19 patients that were clinically well-characterized, with moderate or critical disease, as well as five healthy controls. They identified nine epithelial and 13 immune cell types and performed comprehensive comparisons between patients with critical and moderate COVID-19 and healthy controls. In differential abundance analysis of the cellular landscape, they observed depletion in basal cells and enrichment in neutrophils in critical cases compared with both healthy controls and moderate cases. Additionally, they applied differential expression analysis comparing cells from patients with different disease severity for each cell type and identified transcriptional profiles characterizing patients with critical or moderate disease in these cell types. Specifically, they observed higher expression of some inflammatory mediators in non-resident macrophages (nrMa), and lower levels of some typical anti-viral markers in cytotoxic T cells (CTL) in severe cases compared to moderate cases.

As the results derived in [5] are based on initial clustering into cell types, variable behavior within cell types could be overlooked. To better interpret the differences in immunological responses between patients with critical and moderate disease, we focused on immune cells from samples of these patients (Fig. 4a,b) and applied DA-seq. Five DA cell subpopulations were identified: *DA*1 and *DA*2 are more abundant in critical cases; *DA*3, *DA*4 and *DA*5 are more abundant in moderate cases (Fig. 4c,d, Supplementary Fig. S1c). Subpopulation *DA*3 covers the majority of the monocyte-derived dendritic cell (moDC) cluster. The depletion of moDC in critical cases was also reported in [5]. Other DA subpopulations are sub-clusters within the 13 well separated immune cell types, which have been overlooked in the original clustering-based analysis. To identify the distinct transcriptional profile of these subpopulations, we compared each DA subpopulation to its immediate neighborhood, i.e. complementary cells to the DA subpopulation within the corresponding cluster of known immune cell type. Characterization of these DA subpopulations by gene markers (Fig. 4e) provides important insights on mechanisms associated with COVID-19 disease severity. These DA subpopulations show distinct profiles that separate them from the complementary cells within their corresponding clusters (Fig. 4f), which clustering-based analysis performed in [5] failed to report.

**Figure 4:**
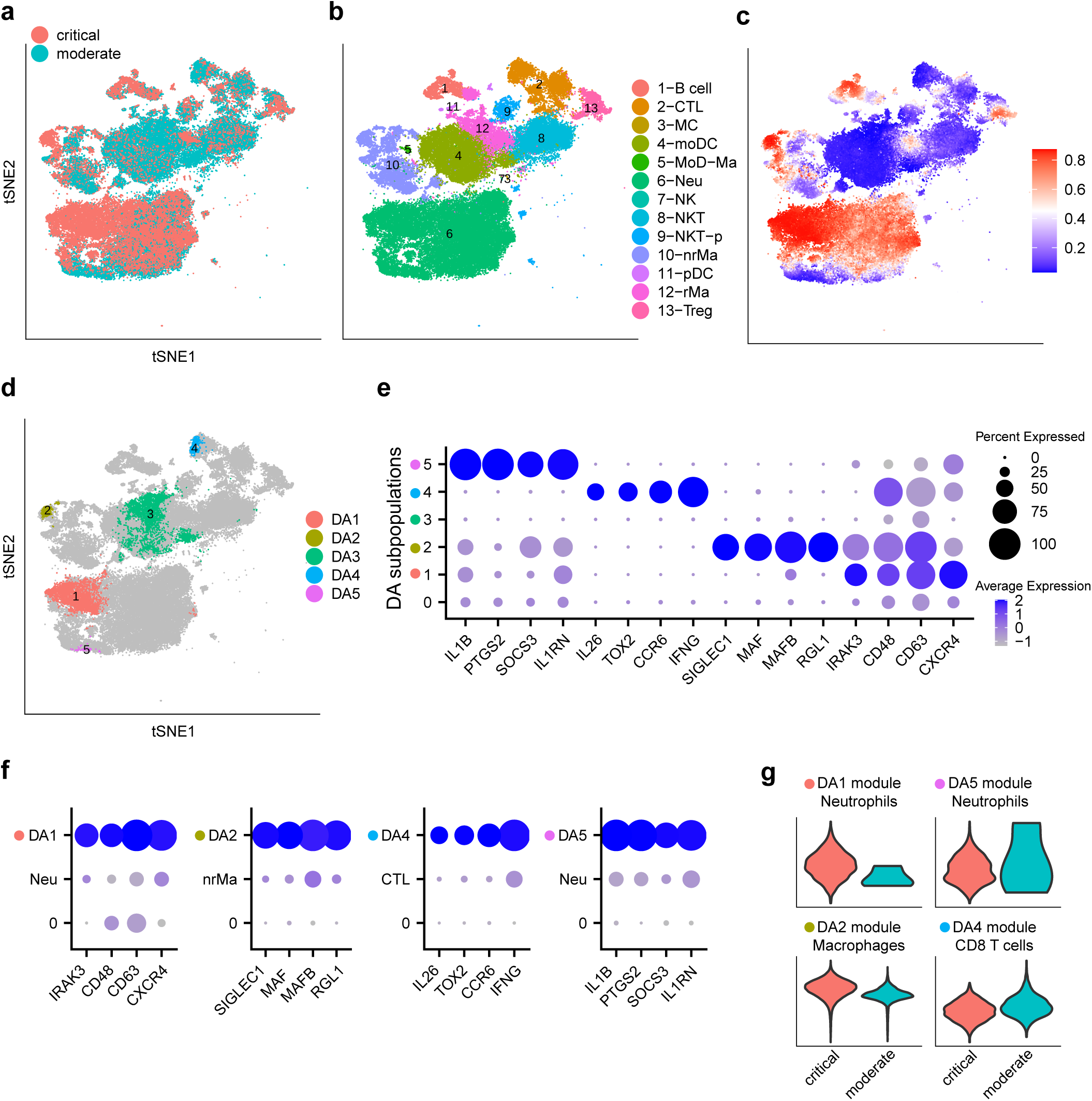
Comparing immune cells from patients with severe and moderate COVID-19. **a-f** Show data from Chua et al. [5]. **a-d** t-SNE embedding of 80,109 cells. **a** Cells colored by disease severity of COVID-19, critical or moderate. **b** Cells colored by cluster labels from [5]. **c** Cells colored by DA measure. Large (small) values indicate a high abundance of cells from the pool of critical (moderate) cases. **d** Five distinct DA subpopulations obtained by clustering cells from the top 5% and bottom 5% quantiles of the DA measure in **c. e** Dot plot for markers characterizing the selected DA subpopulations. Details are as in Fig. 2**e. f** Dot plots for markers of DA subpopulations, comparing each DA subpopulation to the complementary part in the corresponding cluster. **g** Validation on data from Liao et al. [6]. Violin plots comparing gene module scores between critical and moderate cases in matching cell types of data from [6]. Gene modules are defined from DA subpopulations in **d**.

Both cell subpopulations *DA*1 and *DA*5 are within the neutrophil cluster. However, they represent two distinct subsets of neutrophils (Fig. 4e,f, Supplementary Fig. S4a,b). Subpopulation *DA*1 is more abundant in critical cases, and shows elevated expression of activation markers *CD48,CD63* [27, 28]. *Further, expression of another DA*1 marker *CXCR4* has been reported to be associated with acute respiratory distress syndrome (ARDS) [29] and allergic airway inflammation [30]. On the contrary, subpopulation *DA*5 is more abundant in moderate cases, and is characterized by the expression of the inhibitory and anti-inflammatory gene *IL1RN* [31], as well as *SOCS3*, an important regulator in restraining inflammation with previously characterized functions in regulating cytokine signaling and the subsequent response [32, 33, 34]. Another marker enriched in *DA*5 is *PTGS2* (COX2) which has a controversial role and can both promote and constrain inflammation. Enrichment of *PTGS2* expressing neutrophils in moderate patients may suggest its inhibitory role in COVID-19. This provides invaluable insights on the use of nonsteroidal anti-inflammatory drugs (NSAIDs), which is under debate [35]. We note that, while abundances of neutrophils might be affected due to sensitivity to isolation techniques, our differential abundance analysis of neutrophils could still reflect real biological processes.

Subpopulation *DA*2 is a subset of nrMa, and is more abundant in critical cases. Markers of *DA*2 include *RGL1, MAFB* and *SIGLEC1* (Fig. 4e,f, Supplementary Fig. S4c). *RGL1* and *MAFB* are associated with M2 state or alternatively activated macrophages [36, 37]. Interestingly, *MAFB* and *SIGLEC1* have also been reported as a maturation markers of alveolar macrophages [38] and may have implications in mediation of pathology by tissue resident macrophages in COVID-19 lung pathology [6].

Subpopulation *DA*4 is a subset of CTL, and is more abundant in moderate cases. This subpopulation is characterized by high expression of *IFNG* (Fig. 4e,f, Supplementary Fig. S4d). This observation is consistent with the descriptions in [5], where CTLs expressing anti-viral markers were found in patients with moderate COVID-19.

Immunological profiles identified through DA-seq as discussed above should be predictive if they reflect real biological mechanisms in COVID-19 patients. To inspect whether these differential abundance trends are shared in another cohort of COVID-19 patients, we examined a second COVID-19 dataset from [6]. In [6], bronchoalveolar lavage fluid immune cells from COVID-19 patients with different disease severity were sequenced and characterized. To facilitate the analysis, we defined gene signatures (see Methods) that are enriched in our detected DA subpopulations *DA*1, *DA*2, *DA*4 and *DA*5 shown in Fig. 4d. Gene module scores (see Methods) for these gene signatures were computed and compared between COVID-19 patients with moderate and critical disease in matching cell types from the second COVID-19 dataset [6]. Specifically, module scores of gene signatures for *DA*1 and *DA*5 are computed in neutrophils from [6] (of note, there are only four neutrophils out of 7,101 immune cells in moderate cases); module score of gene signature for *DA*2 is computed in macrophages from [6]; module score of gene signature for *DA*4 is computed in CD8 T cells from [6]. The differences between the module score distributions of the critical versus moderate cases (Fig. 4g) are consistent with the enrichment of these signatures within the DA subpopulations in Fig. 4d.

### Additional datasets

In Ximerakis et al. [15], transcriptomes of brain cells from young and old mice are profiled. We applied DA-seq and detected cell subpopulations more abundant in brains from young mice with respect to old mice. Characterizations of these DA subpopulations can be found in Supplementary Note 2 and Supplementary Figs. S5,S6.

In addition, we applied DA-seq to two simulated datasets, and compared the results to Cydar [12]. The first datasets is based on the scRNA-seq data from [7], and the second is a perturbed Gaussian mixture model. A detailed description of both datasets as well as the results can be found in Supplementary Note 3, Supplementary Figs. S7,S8.

## Discussion

In this work, we present DA-seq, a novel multiscale approach for detecting subpopulations of cells that have differential abundance (DA) between scRNA-seq datasets from two biological states. This approach enables us to robustly delineate regions of substantially differential abundance between these two samples. In contrast to existing methods, the subpopulations of cells we discover are not confined to any predefined clusters or cell subtypes. We applied DA-seq to several scRNA-seq datasets and compared its output to results obtained through conventional methods. DA-seq not only recovered results obtained by standard approaches but also revealed striking novel DA subpopulations which informs on cellular function, identifies known and novel genes in DA subpopulations and greatly increases the resolution of cell type identity in different clinical states of disease.

A potential improvement to DA-seq can be achieved by applying a neural network classifier directly on the input features (gene expression profiles or PCA coordinates) without computing the score vector in Step 1. A network architecture for classification of two classes often contains a logistic regression as its last layer. The layers preceding the last layer can then be viewed as feature extractors trained in a supervised way. These features may substitute our hand-crafted, multi-scale score-vector features. We conducted preliminary experiments using the full-neural-network approach. The results were comparable to DA-seq for the simulated datasets but inferior for the real-world datasets. We conjecture that, for our DA problem, the hand-crafted features allow for a better identification of DA cells because these cells are concentrated in two regions in the score-vector space. On the other hand, the landscape of DA cells in the original gene or PCA space is much more complex. However, it is possible that more sophisticated neural network approaches may outperform DA-seq - especially when a larger number of cell measurements is available.

Proper preprocessing of scRNA-seq data is required to obtain reasonable DA results. It is important to recognize that batch effect removal is a necessary preprocessing step for DA-seq in cases where there are noticeable batch effects between samples. Without proper calibration, the DA subpopulations detected by DA-seq may reflect both biological and technical differences between samples. To address this problem in the context of scRNA-seq, multiple batch effect removal methods have been developed [16, 39, 40, 17]. Furthermore, imputation or denoising for scRNA-seq datasets may also improve downstream analysis and lead to a more accurate differential abundance assessment, as cells are positioned more accurately after imputation [41, 42, 43, 44].

In addition to the comparison between two states discussed above, potential applications of DA-seq could be extended to studies comparing multiple biological states, such as time series studies or subjecting a biological system to multiple perturbations. DA-seq can be applied to such multi-state comparisons by considering all pairwise differences in abundance. Alternatively, one can propose a multi-state score vector and replace the binary logistic regression classifier with a multi-class classifier, such as the softmax classifier.

Practitioners often try to detect intra-cluster differentially expressed genes between two states separately for each cluster [16, 7, 15]. If such intra-cluster differentially expressed genes exist, it means that the distributions of cells from these two states are shifted with respect to each other and, hence, represent two adjacent DA subpopulations: one enriched by cells from the first state and the other enriched by cells from the latter. One example is in the comparison between old and young mice shown in Fig. S5. Cluster 21-MG (Fig. S5b) consists of two DA subpopulations, one enriched with cells from old mouse brains, and the other one enriched with cells from young mouse brains. In this case, differentially expressed genes from intra-cluster analysis will be similar to genes that characterize the DA subpopulations with respect to its immediate neighborhood. However, the intra-cluster analysis neither informs us about differential abundance between the states nor is applicable to data with no cluster-like structure.

In many biological systems, cell populations could be heterogeneous in terms of the expression status of certain markers. For instance, breast cancer cells from an ER positive patient do not express ER in all her cancer cells. This status can be measured at the transcriptional or translational level. An application of DA-seq to data generated in a single scRNA-seq experiment to compare her ER(+) or ER(-) cancer cells will enable identification of subpopultions of cancer cells enriched by ER(+) or ER(-) cells and, thus, allow exploration of the biological differences between these two populations (beyond their difference in ER status). Essentially, this approach allows us to use cells generated in a single scRNA-seq experiment and compare cells conditioned on the expression status of a single marker.

In Step 4 of DA-seq, we characterize each DA subpopulation by markers that differentiate it from the remaining cells using our novel neural network embedded feature selection (*l*_0_-based regularization) method. An alternative method to characterize markers associated with DA subpopulations is to link our cells to cells that have been characterized in earlier studies and documented in cell atlases [45]. Unsupervised methods for cell identification can also be utilized to characterize DA subpopulations by overlapping the DA cells and structures revealed by such unsupervised methods [46, 47, 48, 49].

Taken together, DA-seq represents a major advance in the comparative analysis of two distinct biological states. DA-seq has the ability to uncover important, significant and hypothesis driving data which would normally be completely lost within a cloud of transcriptomic data restrained by strict and arbitrary clustering definitions. We envisage DA-seq to be easily integrated into conventional scRNA-seq analysis pipelines and will facilitate major novel findings in all areas of biological investigation.

## Methods

### Problem setup

Our goal is to detect subpopulations of cells that are differentially abundant between two biological states. Let 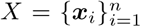 be the gene expression profiles of *n* cells collected in two experiments. In scRNA-seq, the number of genes is typically *∼* 30000, while the number of cells ranges between 10^3^ − 10^6^. Every cell is assigned a binary label *y*_*i*_ *∈* {0, 1}, indicating from which of the two experiments it was collected. We assume that {***x***_*i*_} are independent realizations of an *m* dimensional density function denoted *f*_***x***_(***x***) : ℝ^*m*^ *→* ℝ.

Abstractly, our objective is to detect regions in the gene space where the conditional distribution *f*_***x***|*y*_(***x***|*y* = 1) differs significantly from *f*_***x***|*y*_(***x***|*y* = 0). We say that a point ***x*** is in a differentially abundant (DA) region if

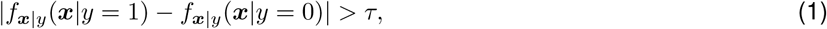

for some threshold *τ*.

One approach to find DA regions is based on *local* two sample tests [50, 51, 52]. A global two sample test determines whether two sets of samples were generated by the same distribution. In contrast, local sample tests also detect the locations of any discrepancies between them. This approach was suggested, among others, by Lun et. al [12] and by Freeman et. al. [50]. These methods compute a test statistic in local neighborhoods around randomly selected cells {***x***_*i*_}. This statistic quantifies the discrepancy between two estimated conditional distributions, 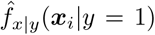 and 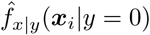, and provides a local *p*-value for each ***x***_*i*_. To correct for multiple testing, the Benjamini-Hoechberg procedure [53] is then applied. Lun et. al. [12] utilized this approach to find DA locations for mass cytometry data.

A different approach for obtaining DA regions was derived by Cazais et. al. [52], where a measure of local discrepancy is computed for all the points in the dataset instead of a random subset. The points with the highest measure of discrepancy are then aggregated into localized clusters in the feature space. Thus, in contrast to the local two sample tests, the output of this approach is a small number of DA regions.

In this work, we derive DA-seq, a new approach for detecting DA regions in scRNA-seq datasets comprising of distinct biological states. DA-seq is based on a multiscale measure of differential abundance computed for each cell. Similarly to [52], our method obtains localized DA regions by clustering the cells with the highest scores. These DA regions represent cell subpopulations with differential abundance between two biological states. Once a DA subpopulation is detected, the next step is to identify characteristic gene expression profiles to distinguish it from other cells. In the following sections, we describe the steps of our approach in detail.

### Step 1: Computing a multiscale score measure

Applying Bayes rule to Eq. (1) gives

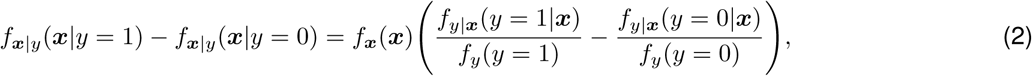

where *f*_***x***_(***x***) is the density function of the cells at a point ***x***. The terms *f*_*y*_(*y* = 1) and *f*_*y*_(*y* = 0) denote the global proportions of the two biological states. Here, state *A* is denoted by *y* = 1 and state *B* is denoted by *y* = 0. The term *f*_*y*|***x***_(*y* = 1|***x***) and *f*_*y*|***x***_(*y* = 0|***x***) are the corresponding *local* proportions around ***x***.

We obtain an estimate of the local proportions via a *k* nearest neighbor estimator, where cell measurement affinities are calculated using the standard Euclidean distance. Note that due to the high dropout rate in scRNA-seq datasets, the Euclidean distance will be taken in the principal component analysis (PCA) or canonical correlation analysis (CCA) space of the gene expression data.

For a specific cell ***x***_*i*_, we define at a scale *k* the *k*-nn estimator of *f*_*y*|***x***_(*y* = 1|***x***_*i*_) and *f*_*y*|***x***_(*y* = 0|***x***_*i*_) by

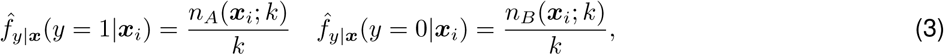

#### Algorithm 1

DA-seq: detecting differential abundance (DA) subpopulations in scRNA-seq

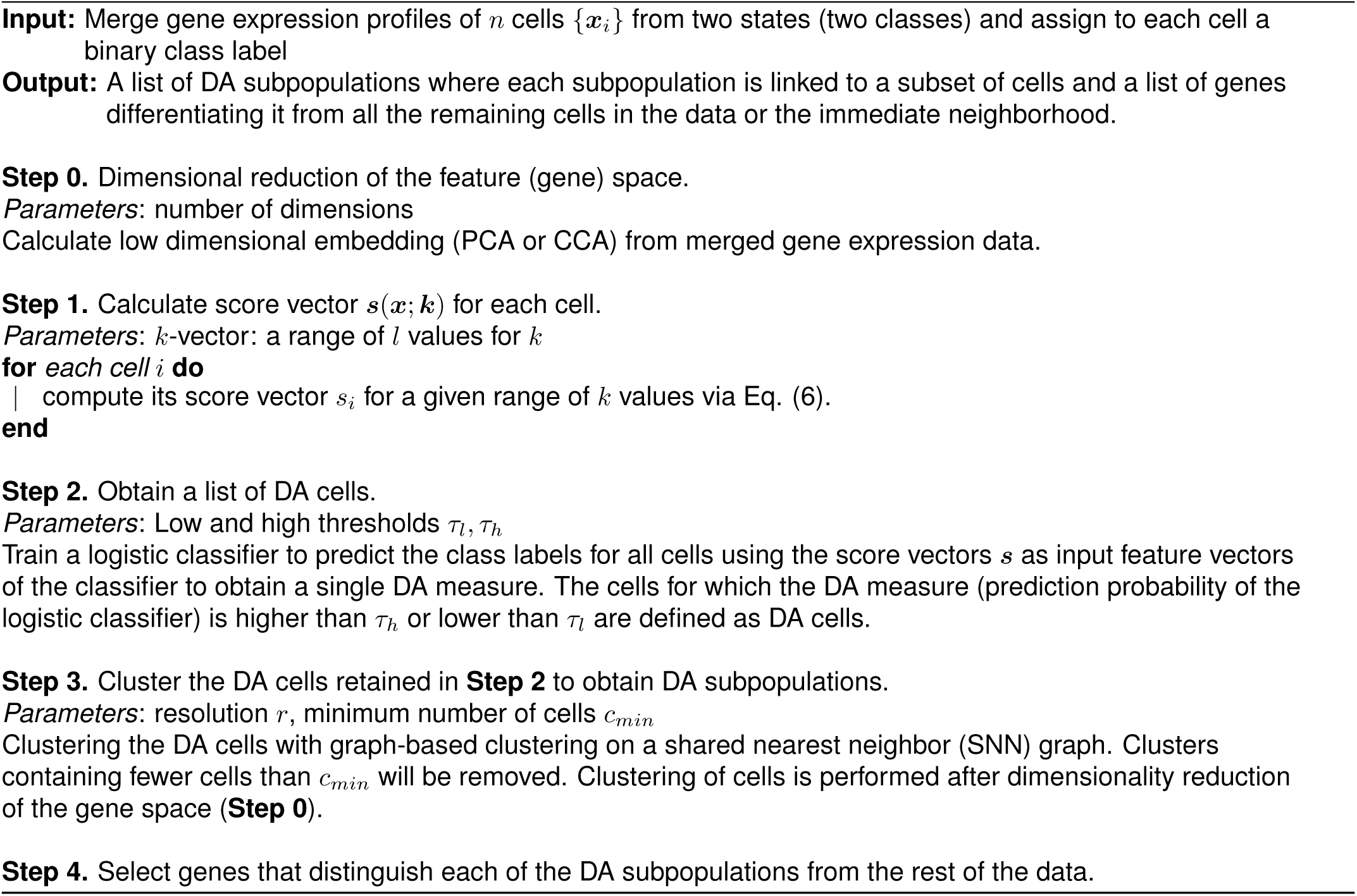

where *n*_*A*_(***x***_*i*_; *k*) and *n*_*B*_(***x***_*i*_; *k*) denote the number of cells among the *k* nearest neighbors of ***x***_*i*_ from biological state *A* and state *B*, respectively. In addition, we define the estimators 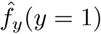 and 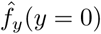 by

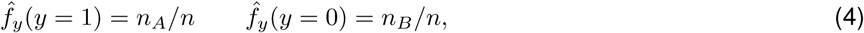

where *n*_*A*_ and *n*_*B*_ are the total number of cells in biological state *A* and *B*, respectively, and *n* = *n*_*A*_ + *n*_*B*_. Inserting (3) and (4) into (2) yields

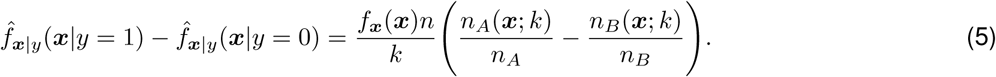

The estimator in Eq. (5) depends on the choice of the number of neighbors *k*. However, a single global value for *k* may be appropriate only for certain regions in the data while being completely suboptimal in other regions. We therefore compute *n*_*A*_(***x***; ***k***) and *n*_*B*_(***x***; ***k***) with a *k*-vector at *l* different nearest neighborhood scales ***k*** = [*k*_1_, *…, k*_*l*_]. We define the normalized score vector ***s***(***x***_*i*_; ***k***) = [*s*_1_(***x***_*i*_; *k*_1_), *…, s*_*l*_(***x***_*i*_; *k*_*l*_)] by

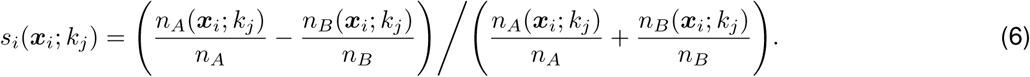

The numerator in (6) is proportional to the *k*NN estimator in (5) while the denominator normalizes the values to be within the range [−1, 1].

Fig. 1 (**Step 1**) illustrates the qualitative behaviour of the score vector ***s***(***k***) for three cells located in different regions of the data. The vector *S*_1_ at the top contains positive entries and corresponds to a cell ***x***_*i*_ in a DA region where *f*_*y*|***x***_(*y* = 1 |***x***_*i*_) *> f*_*y*|***x***_(*y* = 0 |***x***_*i*_). Thus, the score is high for small values of *k*. As *k* increases, the score typically decreases since at this scale the neighbors may contain a more balanced proportion of cells from the two biological states and even include neighbors positioned outside of the DA region.

#### Choice of distance metric

We explored the use of diffusion distance [54] instead of the standard Euclidean metric when calculating the *k*NN estimator in the first simulation data (described in Supplementary Note 3 and Supplementary Fig. S7). Compared with Euclidean distance, there are more outlier cells that lead to a false positive DA subpopulation in results generated with diffusion distance (Supplementary Fig. S9).

#### Computational complexity

The computation of a *k*-nearest neighbor (*k*-nn) graph may be a computational bottleneck for very large datasets. A standard method to compute *k*-nn graphs is via the application of *kd* trees [55]. The complexity of constructing a *kd* tree is *O*(*n* log(*n*)), and the average complexity for finding *k*-nearest neighbors is bounded by *O*(*kn* log(*n*)). For datasets on the order of millions of cells, fast approximate approaches, such as [56, 57], can be applied to increase the scalability of this step.

### Step 2: Computing a DA measure for each cell

The output of Step 1 consists of multiscale score vectors. Cells in DA subpopulations whose neighborhoods are enriched with cells from one biological state tend to be closer to each other in the *l*-dimensional score space than cells whose neighborhoods are enriched with the other biological state or not enriched by any of the states.

Our task in Step 2 is to map the *l* dimensional score vector ***s***(***x***; ***k***), defined in (6), into a single *DA measure* for each cell. To that end, we use a logistic regression classifier. The classifier is trained to predict the class label *y*_*i*_ of each cell given its *l*-dimensional score vector ***s***(***x***_*i*_; ***k***). Specifically, we compute a vector ***w***^*\**^ that minimizes the following loss,

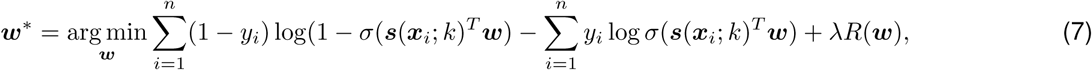

where *σ* is the sigmoid function and *λR*(***w***) is the regularization term. The classifier is trained to increase *σ*(***s***(***x***_*i*_; *k*)^*T*^ ***w***^*\**^) if *y*_*i*_ = 1 and decrease its value if *y*_*i*_ = 0, and thus assigns a numerical value between 0 and 1 based on each cell’s score vector,

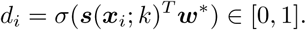

We employ a regularized logistic classifier with ridge penalty by default. The importance of the regularization term is to induce smoothness of the logistic output, such that the cells chosen as DA are localized. In comparison, applying the logistic classifier without regularization produces results with more outliers.

The data is split into *f* folds. For each fold, the model trained on the remainder *f* − 1 folds is used to get the prediction probability. The penalty parameter *λ* for each model is selected by cross-validation. These steps may be repeated for several independent runs and the average is used as the final DA measure for each cell. Notably, the properties of the logistic classifier imply that a high value of *d*_*i*_ is a strong indication that the cell is located in a (score-vector space) region enriched with positive labels, and vice versa.

Fig. 1 (**Step 2**) illustrates the output of the logistic regression classifier. The figure shows a heatmap, where each cell is colored by the prediction probability of the logistic classifier, i.e. its DA measure. The cells that reside in DA regions are determined by thresholding the prediction probability of the logistic classifier. Thus, we determine that ***x***_*i*_ is part of a positive DA region if *d*_*i*_ *> τ*_*h*_ and of a negative DA region if *d*_*i*_ *< τ*_*l*_.

### Step 3: Clustering the DA cells into localized regions

This step involves clustering the subset of cells (DA cells) whose DA measure (prediction probability of the logistic classifier) is above *τ*_*h*_ or below *τ*_*l*_ into localized regions. These DA regions represent cell subpopulations with difference in abundances between biological states. Importantly, the clustering is performed in the original gene space after dimensionality reduction.

We first calculate a shared nearest neighbor (SNN) graph on the whole dataset based on the *k*-nearest-neighbors (*k*NN) graph computed with Euclidean distance with *Seurat* [16, 17] using default parameters. Next, a subgraph comprising of DA cells only is extracted from the full SNN graph. Modularity optimization based clustering algorithm implemented in *Seurat* is applied on this subgraph. For robustness, singletons and very small clusters (containing number of cells fewer than a user defined parameter) will be removed as outliers.

A graph-based clustering approach is used here because of its widespread use in scRNA-seq analysis. We note, however, that other clustering methods can be used for this step. The output of this step is a list of DA subpopulations where each subpopulation is assigned a subset of cells. In our next section, we describe a feature selection approach to identify characteristic genes for each DA subpopulation.

### Step 4: Differential Expression Analysis as a Feature Selection Problem

Differential expression analysis (DEA) and feature selection are related tasks. In DEA, one applies univariate statistical tests to discover biological markers that are typical of a certain state or disease. This approach is typically used for its simplicity and interpretability. Univariate approaches treat each gene individually, however, they ignore multivariate correlations. Feature selection, on the other hand, seeks an interpretable, simplified, and often superior classification model that uses a small number of genes. Here, we use our recently proposed embedded feature selection [13] method to discover for each DA subpopulation a subset of genes that collectively have a profile characteristic for that subpopulation, and thus separates it from the rest of the data.

Given observations 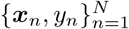, the problem of feature selection could be formulated as an empirical risk minimization

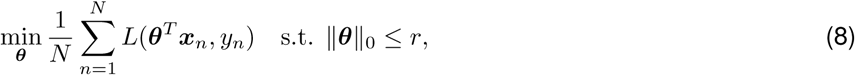

where *r* is the number of selected features, *L* is the loss function and ***θ*** are the parameters of a linear model or more complex neural net model. Due to the *l*_0_ constraint, the problem above is intractable. In practice, the *l*_0_ norm is typically replaced with the *l*_1_ norm, which yields a convex optimization problem as implemented in the popular LASSO optimization approach [58]. Nonetheless, we recently surmounted this obstacle by introducing a stochastic gates (STG) approach to neural networks, which provides a non convex relaxation of the optimization in (8). Each STG is a relaxed Bernouli variable *z*_*d*_, where ℙ(*z*_*d*_ = 1) = *π*_*d*_, *d* = 1, *…, D*, and *D* is the total number of genes. The risk minimization in (8) could be reformulated by gating the variables in ***x*** and minimizing the number of expected active gates. This yields the following objective

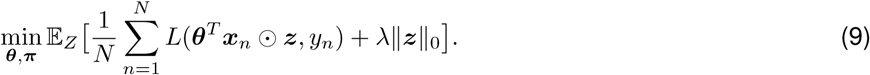

Objective (9) could be solved via gradient descent over the parameters of the model ***θ*** and the gates ***π***. To identify characteristic genes of a DA subpopulation, we train a model that minimizes (9) by sampling multiple balanced batches from the DA subpopulation vs. the backgrounds. Then we explore the distribution of genes that were selected by the model: all *d*’s such that *π*_*d*_ *≥* 0. Note that *λ* is a regularization parameter that controls the number of selected genes; it could be tuned manually for extracting a certain number of genes or, alternatively, using a validation set by finding *λ* which maximizes the generalization accuracy.

An important consideration for tuning *λ* is the potential colinearity between features. Embedded feature selection methods, such as LASSO or STG can capture all correlated features if the regularization parameter is appropriately tuned [13, 59]. In [59], the authors study how correlated variables influence the prediction of LASSO. The authors recommend to decrease the regularization parameter if the correlation between variables is high. A similar behaviour was observed for the *l*_0_ based STG [13].

In this study, STG is used for binary classification (DA subpopulation vs. background cells). We use a standard cross entropy loss in Eq. 9 defined by,

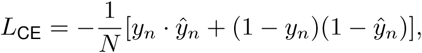

where the predictions *ŷ*_*n*_ and 1− *ŷ*_*n*_ represent the predicted probabilities that the *n*-th cell belongs to the DA sub-population and background, respectively. To obtain a probabilistic interpretation for *ŷ*_*n*_, we use the common sigmoid function

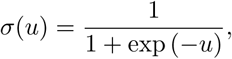

which is in the range of [0, 1]. Using the sigmoid the predicted DA probability is computed as 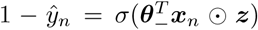. Furthermore, the predicted background probability is computed by 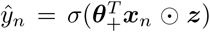, where *θ*_+_ and *θ*_−_ are coefficients for predicting DA and background cells respectively. We then define the STG-score for the experimental section by applying a sigmoid to the difference between the linear predictions of DA and background, that is 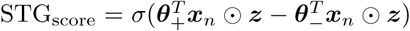. Training of STG is performed using gradient decent with a learning rate of 0.1 using 3000 epochs. These values where observed to perform well across all our experiments.

### Choice of parameters

Parameters for the DA-seq algorithm are outlined in Algorithm 1 under each step.

The choice of the range [*k*_1_, *…, k*_*l*_] in the *k*-vectors should be guided by the data at hand, typically, the lower limit *k*_1_ is the smallest number of cells that a user will consider a meaningful region. The upper limit *k*_*l*_ can be adjusted to the minimal value for which the score, for most cells, converges to the same value. We explored the use of different *k*-vectors on the first simulation data (described in Supplementary Note 3 and Supplementary Fig. S7). The output of Step 2 (selected DA cells by retaining only top and bottom quantiles of the DA measure) for different *k*-vectors but with the same threshold parameters *τ*_*l*_, *τ*_*h*_ is shown in Supplementary Fig. S10a. These results indicate that smaller *k*_1_ (lower limit of the *k*-vector) yielded more outlier cells; whereas the results are less sensitive to the value of *k*_*l*_ (upper limit of the *k*-vector). Given these results, increasing *k*_1_ leads to smoother results in practice.

Another challenge is to determine the threshold parameters *τ*_*l*_, *τ*_*h*_ in Step 2. The ideal solution would be to retain only cells with a neighborhood that is ‘significantly’ DA. A common technique to estimate statistical significance is to perform random permutations and construct the null distribution. We examined the possibility of using permutations in DA-seq for automatic choice of *τ*_*l*_, *τ*_*h*_. To obtain the null distribution, we run the first two steps of DA-seq on randomly permuted cell labels (biological state of each cell), which creates a scenario of uniformly distributed labels without any DA behavior. Let us denote by *p*_*min*_ and *p*_*max*_ the minimum and maximum value of the DA measure (prediction probability of the logistic classifier) on the data with permuted labels. The threshold parameters could be defined as *τ*_*l*_ = *p*_*min*_, *τ*_*h*_ = *p*_*max*_, such that only cells with a DA measure more extreme than random DA measures are retained. In the first simulation data (described in Supplementary Note 3 and Supplementary Fig. S7), we found that using *τ*_*l*_ = *p*_*min*_, *τ*_*h*_ = *p*_*max*_ successfully recovered cells from our artificial DA sites (true positive DA cells) and only introduced very few false positive cells (Supplementary Fig. S10b). However, in real scRNA-seq data from [7], using *τ*_*l*_ = *p*_*min*_, *τ*_*h*_ = *p*_*max*_ yielded results with more than 70% cells retained as significant DA cells (Supplementary Fig. S10c). Thus, considering the complexity and noise in real scRNA-seq datasets, we recommend to set *τ*_*l*_, *τ*_*h*_ based on top and bottom quantiles of the DA measure. If the number of cells from the two biological states are not balanced, as is the case for Fig. 2d on data from Sade-Feldman et al. [7], it is reasonable to use different top and bottom quantiles.

In Supplementary Table S1, we listed parameters used in all datasets presented in the manuscript.

### Preprocessing of scRNA-seq datasets

The R package *Seurat* was used for most preprocessing steps for the scRNA-seq datasets discussed in the paper. Details are described below. In datasets from [5, 7, 15], the preprocessing steps were exactly the same as in the original papers. In the dataset from [14], data integration with *Seurat* [17] was used to remove batch effect, instead of regressing out batch during data scaling. t-distributed stochastic neighbor embedding (t-SNE) embedding for data visualization was calculated with FIt-SNE [60].

#### Melanoma dataset

Transcripts per million (TPM) scRNA-seq data was obtained from [7]. We then performed data scaling and principal component analysis (PCA) with *Seurat*. Following the steps implemented in [7], we calculated the variance for each gene and kept only genes with variance larger than 6 as an input for PCA; the top ten PCs were retained for the calculation of t-SNE embedding and DA analysis.

#### Mouse embryonic dataset

Raw count matrices of scRNA-seq data from two time points E13.5 and E14.5 (two replicates each) were obtained from [14]. For each sample, we used *Seurat* to perform data normalization, scaling, variable gene selection, PCA, clustering and t-SNE embedding calculation. As in [14], markers *Col1a1,Krt10* and *Krt14* were used to select dermal clusters: only cells in clusters with expression of *Col1a1* and no expression of *Krt10,Krt14* were retained for further analysis. After selecting dermal cells, we used *Seurat* data integration to merge data and remove batch effects. PCA was performed on the integrated data, and the top 40 PCs (the same as in [14]) were used to calculate the t-SNE embedding and for DA-seq analysis.

For the five detected DA subpopulations, marker genes were identified using the FindMarkers() *Seurat* function with the ‘negbinom’ method, comparing each DA subpopulation to the rest of the cells. The top 100 genes enriched in the DA subpopulation (or all genes if the number of marker genes is fewer than 100) were selected as a gene signature/module for each DA subpopulation.

For validation, raw count matrices of scRNA-seq data from time points E13 and E15 were downloaded from [26]. For each sample, *Seurat* was used to process the data and generate clusters. Marker gene *Col1a1* was used to select dermal cells. Only dermal cells from both samples were retained and merged for further analysis. The *Seurat* function AddModuleScore() was used to calculate module scores for gene modules of DA subpopulations described above.

#### COVID-19 datasets

The *Seurat* object of integrated data from [5] was downloaded. PCA was performed on the integrated data with 2000 variable features. The top 90 PCs were retained for DA-seq analysis and as input for t-SNE embedding. Immune cells were selected based on cell type labels obtained from the ‘meta.data’ slot of the downloaded object. For detected DA subpopulations *DA*1, *DA*2, *DA*4 and *DA*5, marker genes were identified using the FindMarkers() *Seurat* function with the ‘negbinom’ method, comparing the DA subpopulation to remaining cells in clusters 6-Neu, 10-nrMa, 2-CTL, and 6-Neu, respectively. The top 100 genes enriched in the DA subpopulation (or all genes if the number of marker genes is fewer than 100) were selected as a gene signature/module for each DA subpopulation.

For validation, the *Seurat* object of data from [6] was downloaded. Cell type information was obtained from the ‘meta.data’ slot of the downloaded object. The *Seurat* function AddModuleScore() was used to calculate module scores for gene modules of DA subpopulations described above.

#### Aging brain dataset

The normalized expression matrix of scRNA-seq data from young and old mice was downloaded from Ximerakis et al. [15]. Cell metadata - including cell type, cell sample labels (from young and old mice) - was also obtained from the original paper. As described in [15], PCA was carried out after the identification of variable genes by the “mean variance plot” method from *Seurat*. The top 20 PCs were retained to calculate 2D embedding with t-SNE and as the input for DA-seq.

## Code availability

An R package implementation of DA-seq is freely available at https://github.com/KlugerLab/DAseq. Scripts to reproduce the analysis and figures presented in this manuscript are available at https://github.com/KlugerLab/DAseq-paper.

## Acknowledgments

The authors would like to thank Rihao Qu, Manolis Roulis, Jonathan Levinsohn, Peggy Myung, and Shelli Farhadian for useful discussions and suggestions. This work was supported by NIH grants R01GM131642, UM1 DA051410, 2P50CA121974, R01DK121948, R01GM135928, R01HG008383.

## Notes

### Competing Interest Statement

The authors have declared no competing interest.

### Summary of Updates

In the new version analyzed new COVID-19 dataset. We also updated procedures in steps 2-3. We added independent datasets to validate out findings.

