## Supplemental Information for "Detection of differentially abundant cell subpopulations discriminates biological states in scRNA-seq data"

### Supplementary Information

#### Supplementary Note 1: DA statistics calculation

For each region detected by DA-seq, we compute a DA-score along with a p-value to indicate its statistical significance. The 'DA score' for a region is computed via,

$$DA\ score = \frac{\frac{x(R)}{n_x} - \frac{y(R)}{n_y}}{\frac{x(R)}{n_x} + \frac{y(R)}{n_y}}. \tag{10}$$

where $x(R), y(R)$ denote the number of cells in the DA region $R$ from sample $X$ and $Y$ respectively.

Here we present a method to evaluate the differential abundance of DA regions that is applicable for cases where biological replicates for each state or condition are available. We compute the $p$-value based on the nonparametric Wilcoxon rank sum test to estimate statistical significance. This calculation is only applicable for datasets with two biological states $(X, Y)$ and replicated samples in each biological state: $X_1, \ldots, X_{m_X}, Y_1, \ldots, Y_{m_Y}$, with $m_X, m_Y$ denoting number of replicates for $X, Y$. For each sample $S$, a ratio $r$ of a given DA region $R$ is calculated by,

$$r(S) = \frac{N_{S \cap R}}{N_S}$$

where $N_S$ denotes total number of cells in sample $S$, and $N_{S \cap R}$ denotes number of cells within region $R$ and from sample $S$. Then, we compute the Wilcoxon rank sum test for the vectors $r(X) = \{r(X_1), \ldots, r(X_{m_X})\}$ and $r(Y) = \{r(Y_1), \ldots, r(Y_{m_Y})\}$ to assess whether the difference between them is significant. In cases where $m_X$ and $m_Y$ are small, like in [14] ($m_X = 2, m_Y = 2$), we use a standard two sample $t$-test instead of Wilcoxon. If no biological replicates are available, p-values of DA regions can be computed by other methods such as permutation tests, or partitioning of the data.

**a** DA statistics for data from Sade et al

| Subpopulation | DA-score | $p$-value |
|---|---|---|
| DA1 | 0.973 | 1.05e-06 |
| DA2 | 0.970 | 8.17e-05 |
| DA3 | -0.952 | 1.00e-02 |
| DA4 | -0.981 | 4.04e-02 |
| DA5 | -0.939 | 1.40e-04 |

**b** DA-score for data from Gupta et al

| Subpopulation | E14/E13 | E14.2/E13 | E14/E13.2 | E14.2/E13.2 |
|---|---|---|---|---|
| DA1 | 1.00 | 1.00 | 0.983 | 0.990 |
| DA2 | 0.942 | 0.952 | 1 | 1 |
| DA3 | -1 | -1 | - | - |
| DA4 | -1 | -1 | -1 | -1 |
| DA5 | -1 | -1 | -1 | -1 |

**c** DA statistics for data from Chua et al

| Subpopulation | DA-score | $p$-value |
|---|---|---|
| DA1 | 0.998 | 8.03e-03 |
| DA2 | 1.00 | 1.35e-02 |
| DA3 | -0.956 | 1.56e-04 |
| DA4 | -0.992 | 1.42e-01 |
| DA5 | -1.00 | 3.74e-01 |

**d** DA statistics for data from Ximerakis et al

| Subpopulation | DA-score | $p$-value |
|---|---|---|
| DA1 | -0.692 | 1.86e-03 |
| DA2 | -0.661 | 6.22e-04 |
| DA3 | -0.631 | 2.95e-03 |
| DA4 | -0.501 | 7.36e-03 |
| DA5 | -0.662 | 8.17e-03 |
| DA6 | -0.822 | 7.85e-04 |

Supplementary Figure S1: **Statistics of DA subpopulations detected from real scRNA-seq datasets. a** Data from Sade-Feldman et al. [7]. A DA-score $> 0$ ($< 0$) indicates the DA subpopulation is more abundant in samples from non-responders (responders). **b** Data from Gupta et al. [14]. A DA-score $> 0$ ($< 0$) indicates the DA subpopulation is more abundant in samples from E14.5 (E13.5). **c** Data from Chua et al. [5]. A DA-score $> 0$ ($< 0$) indicates the DA subpopulation is more abundant in samples from severe (moderate) patients. **d** Data from Ximerakis et al. [15]. A DA-score $< 0$ indicates the DA subpopulation is more abundant in samples from young mice.

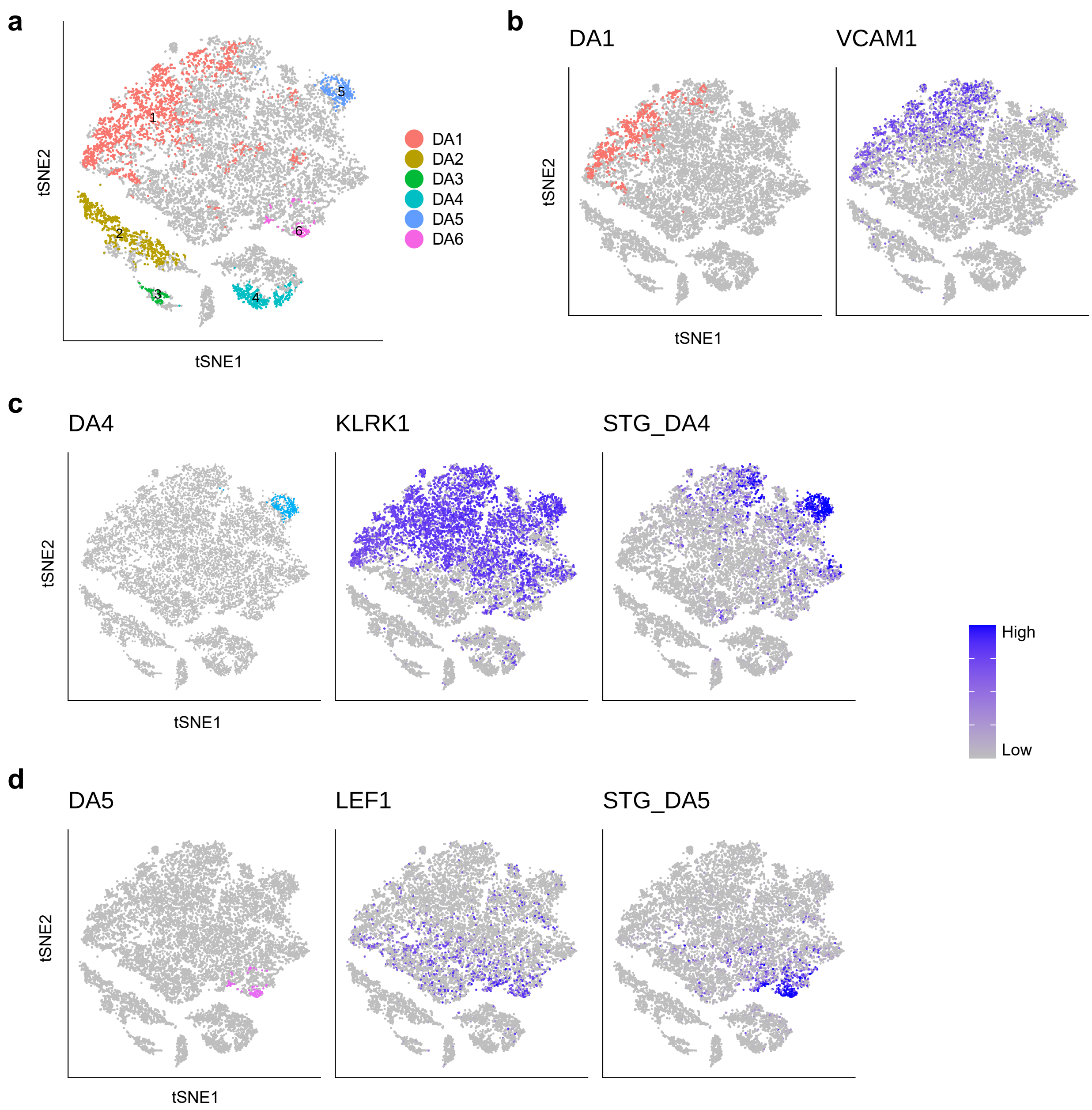


Supplementary Figure S2: **Characterizing DA subpopulations in data from Sade-Feldman et al. [7].** **a** Identified DA subpopulations after relaxing the threshold $\tau_h$ to $20\%$. Subpopulation $DA3$ corresponds to the dendritic cell cluster $G4$. **b-d** Marker gene expression and STG prediction score overlay on t-SNE embedding for DA subpopulations. **b** DA1. **c** DA4. **d** DA5.

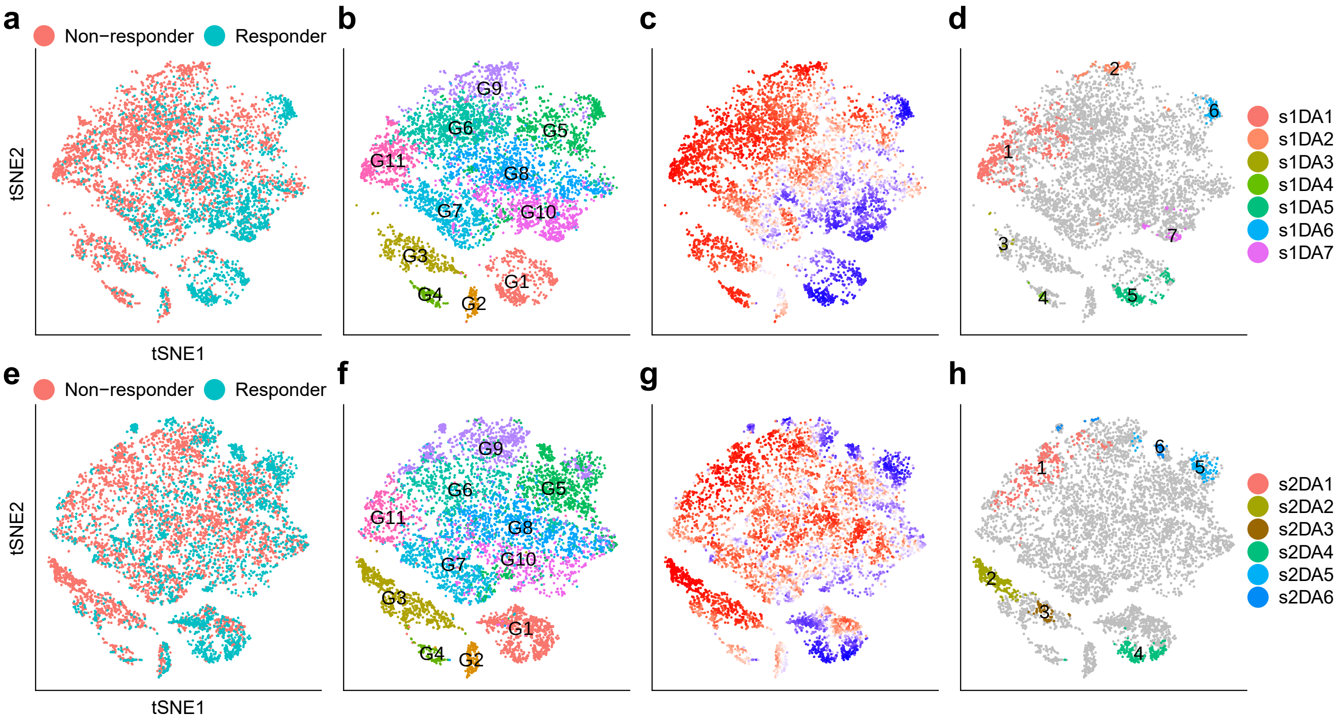


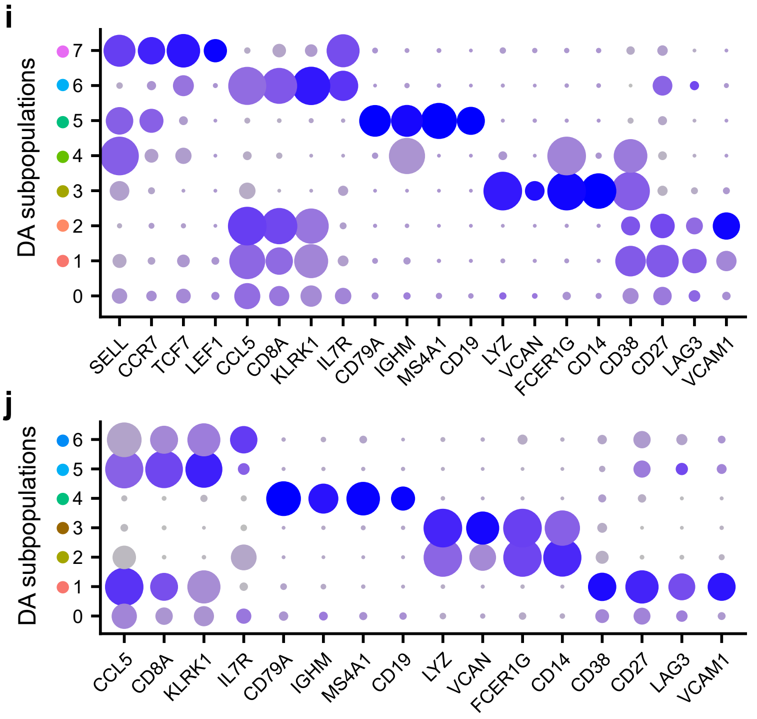


**k**

| Full dataset | Split data1 | Split data2 |
|---|---|---|
| DA1 | s1DA1<br>s1DA2 | s2DA1 |
| DA2 | s1DA3 | s2DA2<br>s2DA3 |
| DA3 | s1DA5 | s2DA4 |
| DA4 | s1DA6 | s2DA5<br>s2DA6 |
| DA5 | s1DA7 | |

Supplementary Figure S3: **Cross-validation on data from Sade-Feldman et al. [7].** Data was split randomly into two sets, each containing half non-responder samples and half responder samples. **a-d** Show results from split dataset 1 on t-SNE embedding of 7,679 cells. **e-h** Show results from split dataset 2 on t-SNE embedding of 8,612 cells. **a,e** Status of response to immune therapy for each cell. **b,f** Cells colored by cluster labels from [7]. **c,g** Cells colored by DA measure. Large/red (small/blue) values indicate a high abundance of cells from the pool of non-responder (responder) samples. **d,h** distinct DA subpopulations obtained by clustering cells from the top $10\%$ and bottom $5\%$ quantiles of the DA measure in **c,g**. **i,j** Dot plots showing marker genes of DA subpopulations. **k** Matched DA subpopulations in the full dataset, split dataset 1 and split dataset 2.

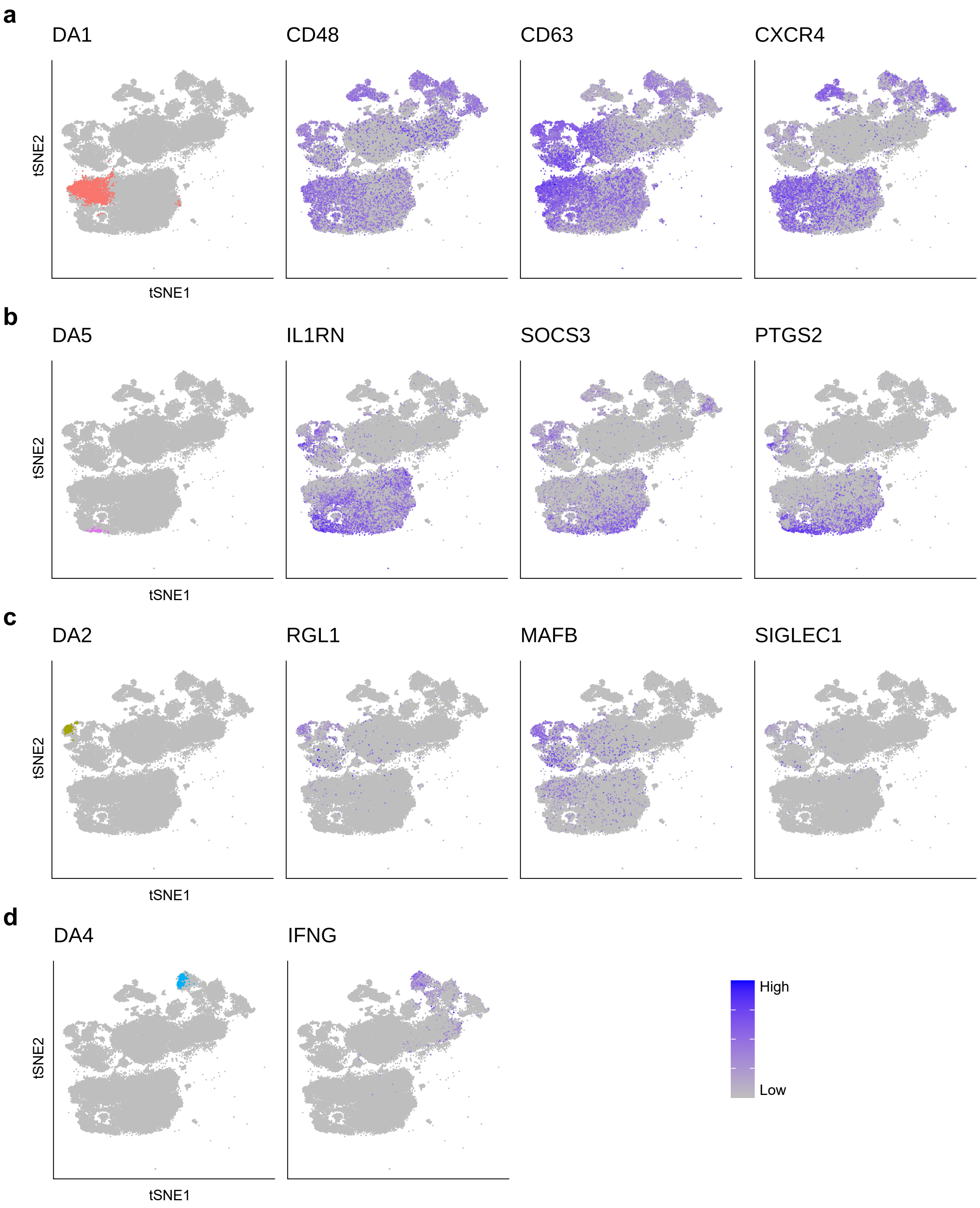


Supplementary Figure S4: **Characterizing DA subpopulations in data from Chua et al. [5].** Marker gene expression overlay on t-SNE embedding for DA subpopulations. **a** DA1. **b** DA5. **c** DA2. **d** DA4.

### Supplementary Note 2: Comparing brain transcriptomic profiles of young and old mice

Ximerakis et al. [15] characterized differences between brain cells of young and old mice (Fig. S5a). As shown in Fig. S5b, they detected 25 distinct cell populations, of which oligodendrocyte precursor cells (1-OPC), neuronal restricted precursors (7-NRP) and immature neurons (8-ImmN) exhibited statistically significant decreases on abundance in old mice.

We applied DA-seq on the merged data and identified six DA subpopulations that are more abundant in young mice, as shown in Fig. S5c,d. DA clusters reported in [15] were identified by DA-seq, specifically, $DA2$ corresponds to 8-ImmN, $DA3$ to 1-OPC, $DA4$ to 7-NRP. Characteristic markers for these DA subpopulations obtained by DA-seq, shown in Fig. S5e agree with markers of the corresponding clusters. In addition, DA-seq identified other DA cell subpopulations. Cells in $DA5$ are classified as oligodendrocytes (2-OLG) but had a different transcriptomic profile: the top marker *Rassf10* (Fig. S5e and Supplementary Fig. S6a) was shown to be associated with neural progenitor cell proliferation and differentiation [61]. Subpopulation $DA6$ is a small subset of 1-OPC and are also characterized by cell cycle related markers, which may represent a proliferative cell subset that is more abundant in brains from young mice. Subpopulation $DA1$ overlaps with a proportion of the microglia (21-MG) cluster and is characterized by the microglia marker gene *Tmem119*.

It is instructive to study the differences between DA cell subpopulations and the clusters that overlap with them. For instance, we examine below how $DA1$ differs from the remaining cells in the microglia cluster (marked as $DA1c$ in Fig. S5d subpanel). The *Tmem119* gene differentiates $DA1$ subpopulation from cells in the rest of the clusters but does not distinguish it from cells from $DA1c$ (Supplementary Fig. S6b). To better identify marker genes that characterize $DA1$, we applied STG to differentiate cells from $DA1$ and cells from $DA1c$. STG identified a total of 71 genes that as a whole differentiate cells in $DA1$ from cells in $DA1c$. Importantly, the gene combination inferred by STG provides a cleaner way to differentiate between the cell subpopulations; this cannot be achieved by individual gene markers (Supplementary Fig. S6c). In Fig. S5f, we show the prediction score of STG based on the combination of these 71 genes. This specific combination of genes represents a particular gene set that may play an important role in microglia during aging.

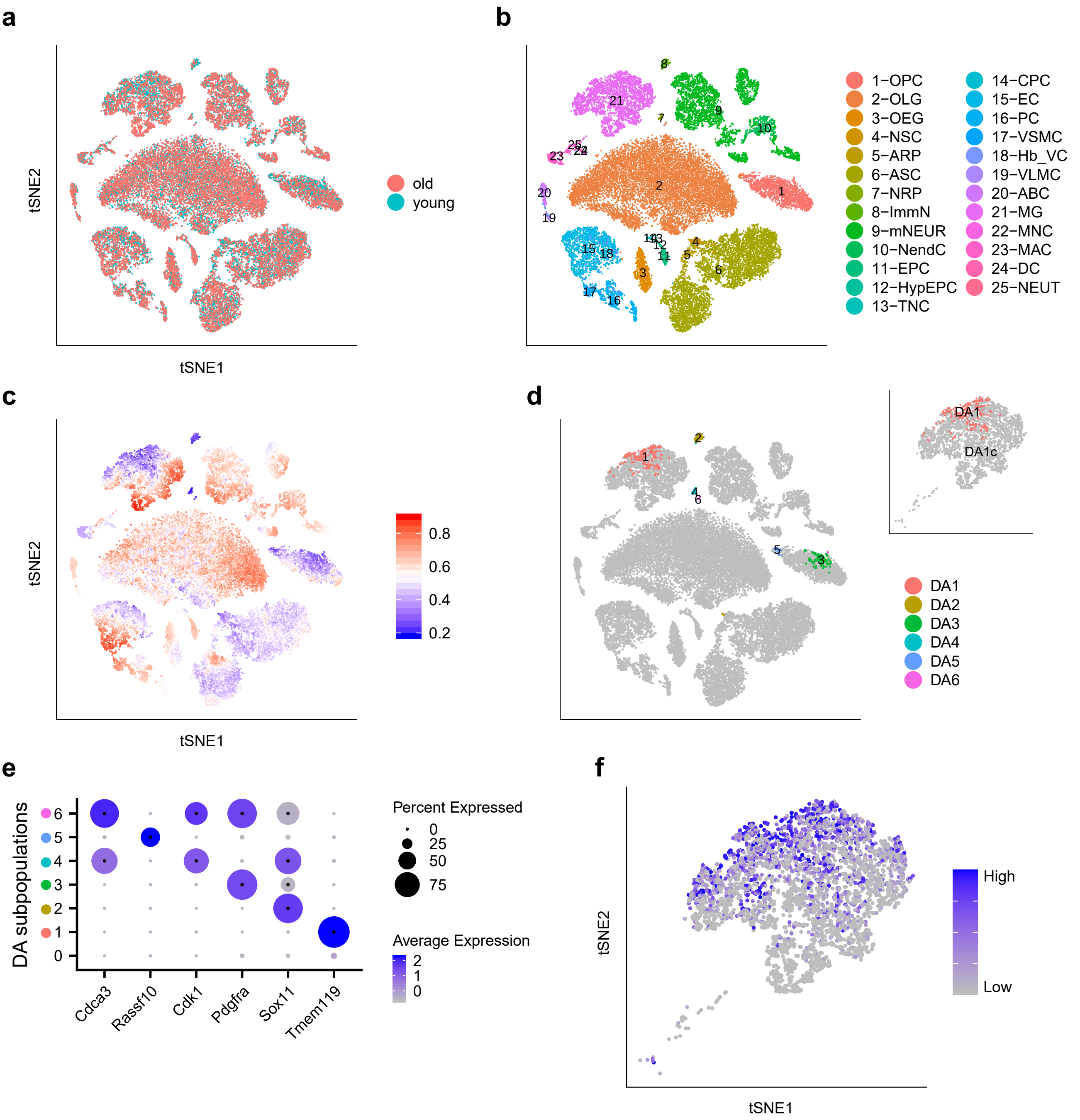


Supplementary Figure S5: **Comparing brain transcriptomics of young and old mice. a-d** t-SNE embedding of 37,069 cells. **a** Cells measured in samples from young mice or old mice. **b** Cluster labels from [15] for each cell. **c** Cells colored by DA measure (prediction probability of the logistic classifier). Large (small) values indicate a high abundance of cells from old (young) mice. **d** Distinct DA subpopulations obtained by clustering cells from the bottom $2\%$ quantile of the DA measure in **c**. **e** Dot plot for markers characterizing the selected DA subpopulations; details are as in Fig. 2**e**. **f** Microglial (cluster 21-MG) cells colored by STG prediction trained to differentiate subpopulation $DA1$ from the remaining microglial cells ($DA1c$ in subpanel of **d**).

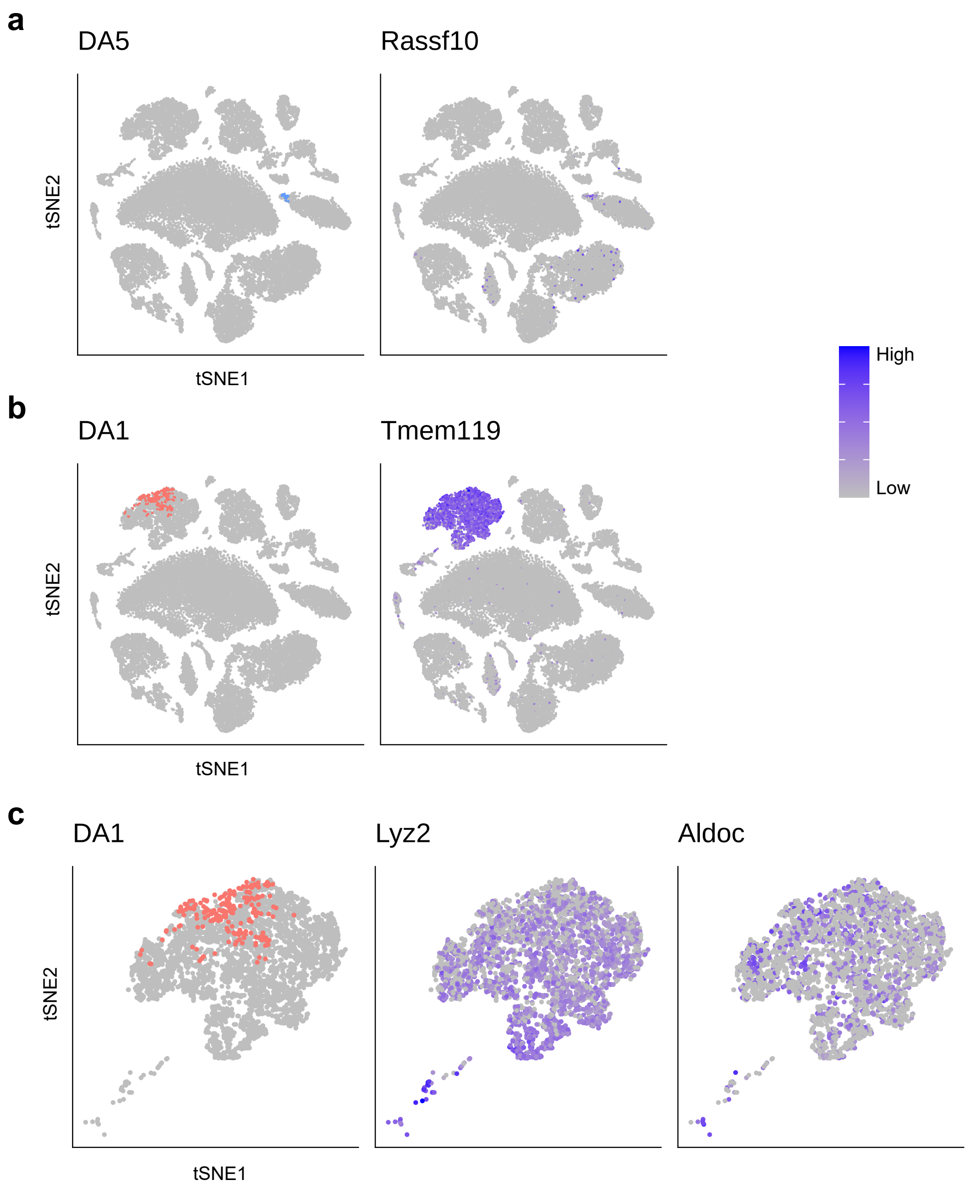


Supplementary Figure S6: **Characterizing DA subpopulations in data from Ximerakis et al. [15].** Gene expression overlay on t-SNE embedding. **a** Marker gene *Rassf10* characterizing subpopulation DA5. **b** Marker gene *Tmem119* differentiating subpopulation DA1 from cells from the rest of the data. **c** Marker genes *Lyz2, Aldoc* identified with standard differential expression analysis to separate subpopulation DA1 from its immediate neighborhood in the same cluster.

### Supplementary Note 3: Simulated datasets

In order to test our algorithm with "ground truth" DA subpopulations, we generated two simulated datasets. The first dataset is based on the scRNA-seq data from [7], and the second on a Gaussian mixture model.

In the first simulation dataset, we used the gene expression profiles from the real data, but assigned labels to cells manually to create artificial DA subpopulations. We selected at random four DA subpopulations based on the $k$NN graph of the data, as shown in Fig S7a. For cells within the DA subpopulations, imbalanced cells labels were assigned at random, with $90\%$ condition $0$, $10\%$ condition $1$, or vice versa; labels for cells outside the DA subpopulations were randomly assigned with $50\%$ condition $0$ and $50\%$ condition $1$ (Fig. S7b). Next, We applied DA-seq to the simulated dataset. The DA measure (prediction probability of the logistic classifier) and final DA subpopulations from our algorithm are shown in Fig. S7c and d, respectively. As can be seen, we successfully recovered all four artificial DA sites and did not introduce false positive subpopulations.

#### Comparison with Cydar

For comparison, we applied Cydar [12] to the simulated dataset. Briefly, Cydar allocates cells into hyperspheres by randomly selecting a proportion of cells as centers and using a fixed radius. Next, Cydar tests for differential abundance in each hypersphere with a negative binomial model. In Fig. S7e, we plotted all hyperspheres obtained from Cydar as circles, positioned at their centers, and colored them based on the magnitude of DA in terms of log fold change within each hyperspheres. The number of cells in each hypersphere is indicated by the size of each circle. Since it is hard to evaluate Cydar's performance with just the hyperspheres, we then merged all significant hyperspheres (with $FDR < 0.05$) and highlighted the cells in Fig. S7f. Clearly, Cydar failed to recover all four DA sites, and identified several false positive sites. Results of Cydar showed limitation of the method on scRNA-seq data due to: 1) centers of hyperspheres are randomly selected, and sometimes can not cover the whole data; 2) radius of the hypersphere is fixed which typically does not match with the actual size of the DA region. In contrast, our multi-scale approach detects the size of neighborhoods with significant differential abundance.

#### Gaussian mixture simulation

We generated the second simulation dataset according to a Gaussian mixture model. Let $\mu_1, \mu_2$ be 10 dimensional vectors, such that each contains two non zero elements (for instance, elements 1-2 for $\mu_1$ and 3-4 for $\mu_2$). The 'cells' were generated according to two distribution functions, $f(\boldsymbol{x}|y = A)$ and $f(\boldsymbol{x}|y = B)$ were $A$ and $B$ correspond to two biological states (Supplementary Fig. S8a)

$$f(\boldsymbol{x}|y=A) = \begin{cases} \mathcal{N}(0,I) & \text{w.p. } 0.9 \\ \mathcal{N}(\mu_1,I) & \text{w.p. } 0.09 \\ \mathcal{N}(\mu_2,I) & \text{w.p. } 0.01 \end{cases} \qquad f(\boldsymbol{x}|y=B) = \begin{cases} \mathcal{N}(0,I) & \text{w.p. } 0.9 \\ \mathcal{N}(\mu_1,I) & \text{w.p. } 0.01 \\ \mathcal{N}(\mu_2,I) & \text{w.p. } 0.09 \end{cases}$$

where $\mathcal{N}(0, I)$ denotes a 10 dimensional standard Gaussian distribution. Thus, the first two features are differentially expressed, and form a DA subpopulation with high abundance of cells from condition $A$. Similarly, the other two differentially expressed features form a second DA subpopulation with high abundance of cells from condition $B$. These two artificial DA subpopulations are highlighted in Supplementary Fig. S8b. Supplementary Fig. S8c shows the output of the logistic classifier trained on the score vector, and Supplementary Fig. S8d shows detected DA subpopulations after clustering cells in the top and bottom $5\%$ quantiles. Applying feature selection through STG, we were able to recover all the differentiating features for this dataset (Supplementary Fig. S8e).

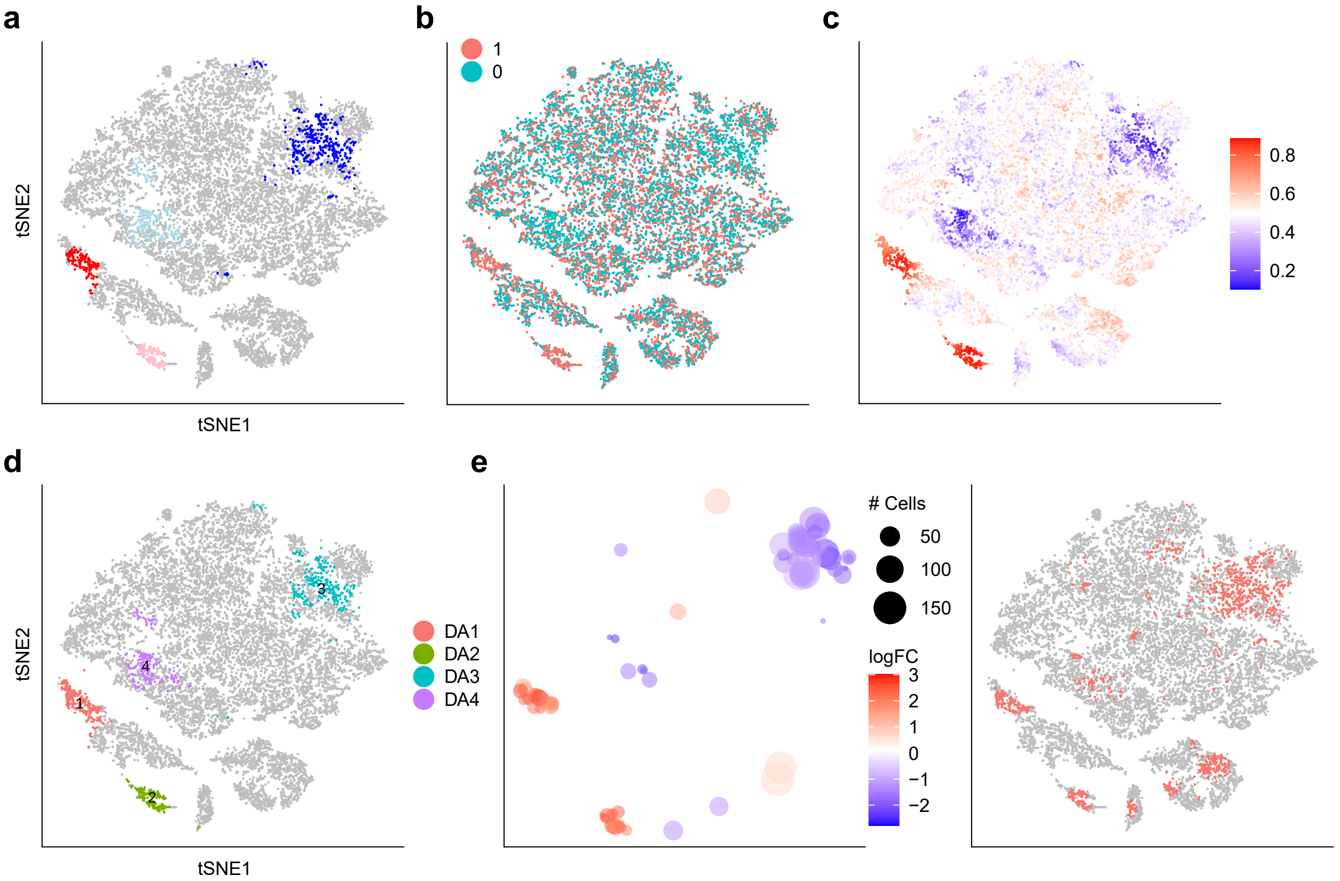


Supplementary Figure S7: **DA results on simulated dataset. a-d** t-SNE embedding of 16,291 cells. **a** Colored cells are four artificial DA subpopulations: two colored with blue hues have more cells from condition 0, while two colored with red hues have more cells from condition 1. **b** Artificial labels for each cell, either condition 0 or condition 1. **c** Cells colored by DA measure. Large (small) values indicate a high abundance of cells from condition 1 (condition 0). **d** Distinct DA subpopulations obtained by clustering cells from the top and bottom $4\%$ quantiles of the DA measure in **c**. **e** Cydar results. Left panel: hyperspheres generated from Cydar, positioned at the center cell of each hypersphere, colored with log fold change of the abundance difference, with size corresponding to the number of cells of each hypersphere. Right panel: cells from significant hyperspheres ($FDR < 0.05$) are highlighted.

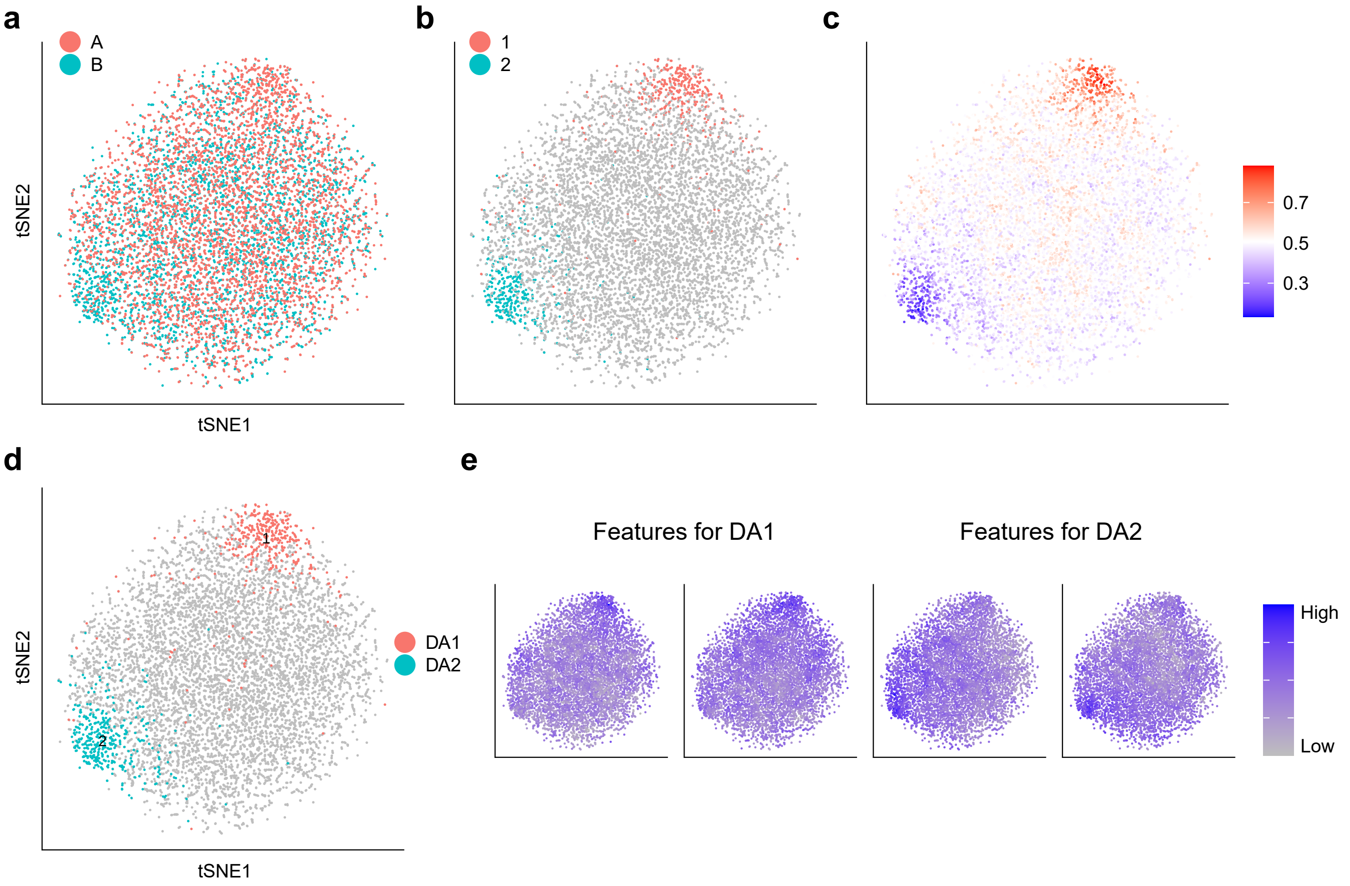


Supplementary Figure S8: **A Gaussian mixture model. a-d** t-SNE embedding of 10,000 cells. **a** Cells colored by its biological condition, A or B. **b** Two artificial DA sites are highlighted. **c** Cells colored by DA measure. Large (small) values indicate a high abundance of cells from condition A (condition B). **d** Distinct DA subpopulations obtained by clustering cells from the top and bottom $5\%$ quantiles of the DA measure in **c**. **e** Cells colored by features selected by STG that differentiate the DA subpopulations.

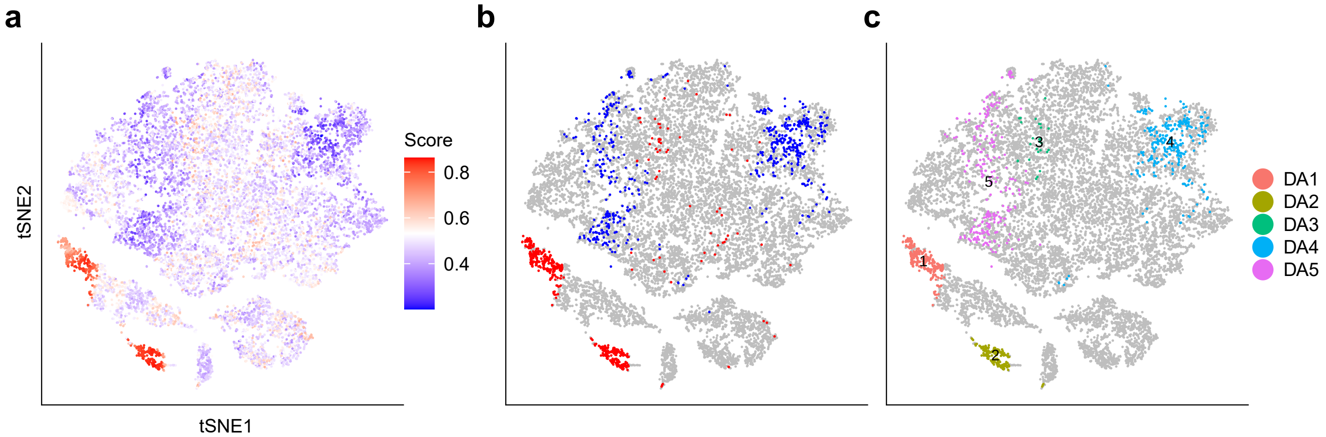


Supplementary Figure S9: **DA-seq with diffusion distance.** t-SNE embedding of 16,291 cells. **a** Cells colored by the DA measure. **b** Highlighted DA cells retaining the top and bottom $4\%$ of values in **a**. **c** Cells colored by distinct DA subpopulations.

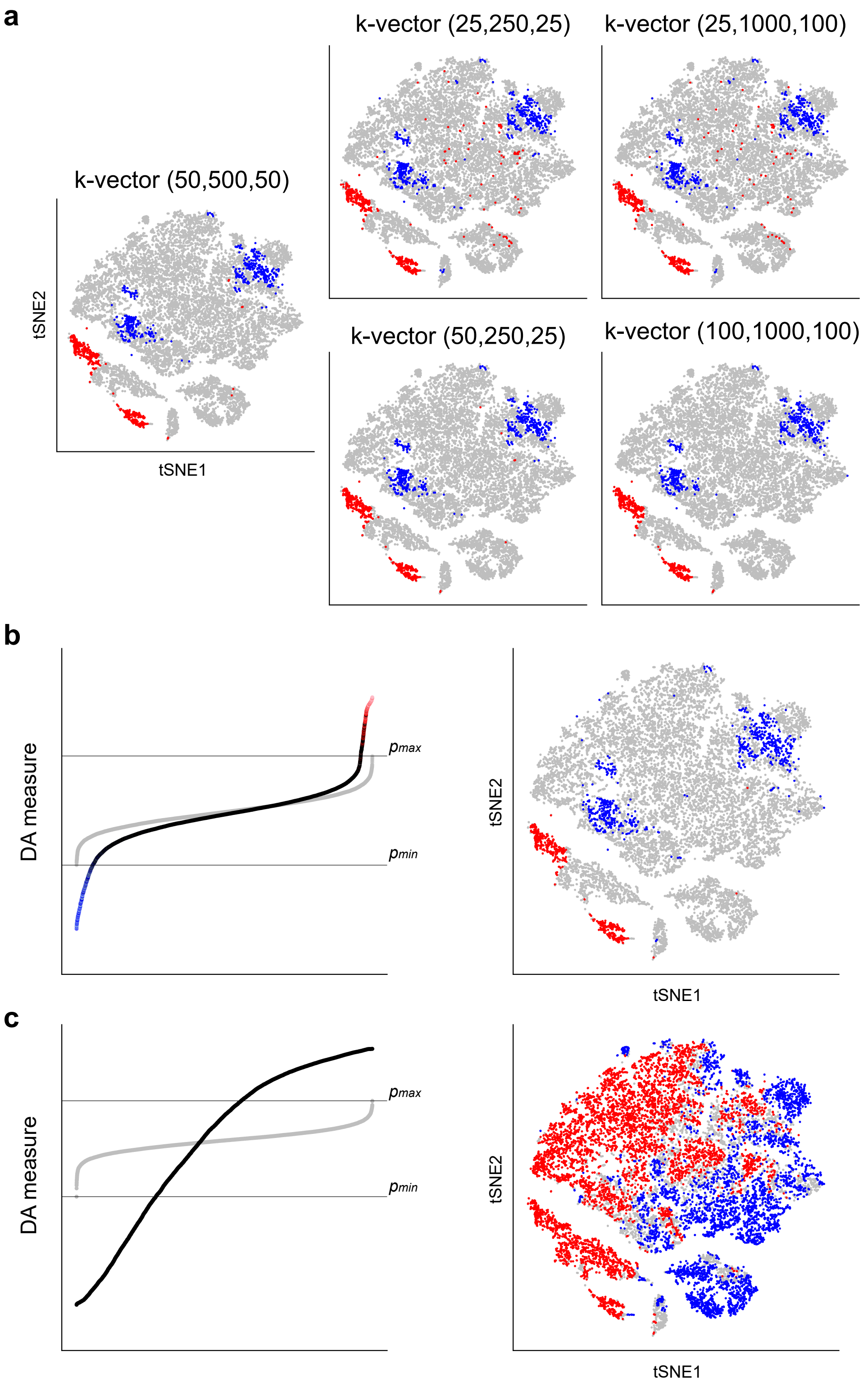


Supplementary Figure S10: **Choice of parameters in DA-seq.** **a** Retained DA cells on the first simulation dataset with different $k$-vectors and identical $\tau_l, \tau_h$. $k$-vector for each subpanel marked in the format of ($k_1, k_l$,steps). **b** Results on the first simulation dataset. Left panel: DA measure (prediction probability of the logistic classifier) for every cell, ordered monotonically, on real labels (black line, cells from artificial DA sites highlighted in red or blue) and permuted labels (gray line). Right panel: retained DA cells with $\tau_l = \min(p_r), \tau_h = \max(p_r)$, $\min(p_r)$ and $\max(p_r)$ marked with two horizontal lines. **c** Results on real scRNA-seq dataset from [7]. Two panels structured similar to **b**.

| **Dataset** | **# cells in dataset** | **$k$-vector** $(\boldsymbol{k_1}, \boldsymbol{k_l}, \boldsymbol{step})$ | **threshold** $(\boldsymbol{\tau_l}, \boldsymbol{\tau_h})$ (*quantile*) | **resolution** $(\boldsymbol{r})$ | **min # cells** $(\boldsymbol{c_{min}})$ |
|---|---|---|---|---|---|
| Sade et al. | 16,291 | (50,500,50) | (0.05,0.9) | 0.01 | 15 |
| Sade et al. split 1 | 7,679 | (50,250,50) | (0.05,0.9) | 0.01 | 10 |
| Sade et al. split 2 | 8,612 | (50,250,50) | (0.05,0.9) | 0.01 | 10 |
| Gupta et al. | 15,325 | (50,500,50) | (0.03,0.97) | 0.05 | 15 |
| Chua et al. | 80,109 | (200,3200,200) | (0.05,0.95) | 0.01 | 15 |
| Ximerakis et al. | 37,069 | (200,1000,100) | (0.02,1) | 0.1 | 15 |
| Simulation | 16,291 | (50,500,50) | (0.04,0.96) | 0.05 | 15 |
| Gaussian mixture | 10,000 | (100,500,50) | (0.05,0.95) | 0.05 | 15 |

Table S1: Parameters used in different datasets.
